# Dystonia-associated Torsins sustain CLCC1 function to promote membrane fusion of the nuclear envelope for NPC biogenesis

**DOI:** 10.1101/2025.11.07.687155

**Authors:** Daria Maslennikova, Harry Baird, Alessia Loffreda, Kaustubh Ramachandran, Ashutosh Kumar, Jelmi uit de Bos, Caroline Ashiono, Katharina Frischer-Ordu, Naemi Luithle, Federico Uliana, Gautam Dey, Stefano Vanni, Madhav Jagannathan, Ulrike Kutay

## Abstract

DYT1 early-onset dystonia is a severe, incurable disorder of the central nervous system caused by mutations in the gene encoding Torsin1A (Tor1A, DYT1). Torsins are ER-resident AAA^+^-ATPases implicated in lipid metabolism, nuclear pore complex (NPC) biogenesis, and lipoprotein secretion, yet their molecular function that underlies the disease pathology has remained incompletely understood. Here, we have utilized *Drosophila melanogaster* and human somatic cells as experimental models to shed light on their mode-of-action. Fly germ cells lacking dTorsin are arrested in development and display defects in the final steps of NPC biogenesis due to a failure in fusion of the inner and outer nuclear membranes. Using proximity labelling of Torsin1A in human cells, we identify the conserved membrane protein chloride channel CLIC-like protein 1 (CLCC1) as a novel Torsin binding partner. Absence of human CLCC1 or its *Drosophila* homolog dClcc1 phenocopied the membrane fusion defects at NPC assembly sites observed upon Torsin deletion. Furthermore, CLCC1 is enriched at arrested fusion sites, suggesting it to be a candidate for the elusive NE membrane fusogen. Importantly, CLCC1/dClcc1 overexpression is sufficient to rescue NPC biogenesis and developmental defects associated with Torsin-loss-of-function. Taken together, our data suggest that Torsin-regulated CLCC1 activity drives membrane fusion during NPC biogenesis and reveal that modulating CLCC1 expression is a promising therapeutic prospect for DYT1 dystonia.

## Introduction

DYT1 early-onset torsion dystonia, caused by mutations in the gene encoding Torsin1A (Tor1A, DYT1), is a severe movement disorder associated with involuntary, repetitive muscle contractions and aberrant body postures (Ozelius et al., 1997). Torsins are AAA+-ATPases of the endoplasmic reticulum (ER) lumen and contiguous perinuclear space that are found in most metazoans but are absent in plants and fungi (Kustedjo et al., 2000, Rose et al., 2015). They show similarity to substrate-threading bacterial Clp AAA+-ATPases (Ozelius et al., 1999). However, unlike these conventional AAA+-ATPases that exploit ATP hydrolysis between neighbouring subunits for substrate threading through a central pore (Hanson & Whiteheart, 2005), Torsins lack the required substrate-binding pore loops and use a distinct mechanism for activation of ATP hydrolysis. Instead, Torsins rely on specialized partner proteins to induce ATPase activity (Brown et al., 2014, Goodchild & Dauer, 2005, Sosa et al., 2014, Zhao et al., 2013). In human cells, these activators are the transmembrane proteins LULL1 and LAP1 which reside in the ER and nuclear envelope (NE), respectively (Fig. S1A). Both harbour a conserved luminal Torsin homology domain that contains a decisive arginine residue to complete the active site of Torsins for induction of ATP hydrolysis (Brown et al., 2014, Demircioglu et al., 2016, Sosa et al., 2014). To date, the identities of potential Torsin substrates that explain its loss-of-function and disease phenotypes are unknown.

Humans express five Torsins (Tor1A, Tor1B, Tor2A, Tor3A and Tor4A) of which Tor1A, the disease-causing founding family member, is the predominant isoform in the developing brain (Jungwirth et al., 2010, Tanabe et al., 2016) (Fig. S1A). Tor1A and Tor1B share 85% similarity and function largely redundantly in other tissues (Jungwirth et al., 2010). In DYT1 early-onset dystonia, deletion of one of two consecutive glutamate residues (ΔE302/303) in the ATPase domain of Torsin1A prevents the binding of Torsin activators LAP1 and LULL1 and thereby ATPase activation (Demircioglu et al., 2016, Naismith et al., 2009, Zhao et al., 2013, Zhu et al., 2010). Consistently, mutations in the *TOR1AIP1* gene, encoding for LAP1, have also been linked to neurological disease phenotypes and primary dystonia (Dorboz et al., 2014, Fichtman et al., 2019).

Although Torsins have been implicated in a variety of cellular processes including NPC biogenesis (Jacquemyn et al., 2021, Kim et al., 2024, Laudermilch et al., 2016, Pappas et al., 2018, Rampello et al., 2020), LINC complex remodelling (Dominguez Gonzalez et al., 2018, Nery et al., 2008), lipid metabolism (Cascalho et al., 2020, Grillet et al., 2016, Hernandez-Ono et al., 2024, Jacquemyn et al., 2021, Shin et al., 2019), ER protein secretion (Hewett et al., 2007, Shin et al., 2019, Torres et al., 2004), as well as vesicular egress of large RNPs and viruses across the NE (Gyorgy et al., 2018, Holper et al., 2020, Jokhi et al., 2013, Maric et al., 2011, Sule et al., 2024), their molecular substrates and mode-of-action have remained incompletely understood. On a cellular level, across a wide range of experimental systems, Torsin loss-of-function leads to the accumulation of evaginations of the inner nuclear membrane (INM) into the perinuclear space (Goodchild et al., 2005, Jacquemyn et al., 2021, Kim et al., 2010, Speese et al., 2012, VanGompel et al., 2015) (Fig. 1A). These NE herniations, or ‘NE blebs’, contain immature NPCs at their collared necks and accumulate ubiquitin and chaperones in their lumen (Kuiper et al., 2022, Laudermilch et al., 2016, Liang et al., 2014, Prophet et al., 2022, Rampello et al., 2020). It has been suggested that these herniations arise by a failure in fusion of the INM and outer nuclear membrane (ONM) during final steps of NPC biogenesis (Rampello et al., 2020), but how this fusion reaction is mediated has remained a longstanding enigma in eukaryotic cell biology (Rothballer & Kutay, 2013).

**Figure 1:**
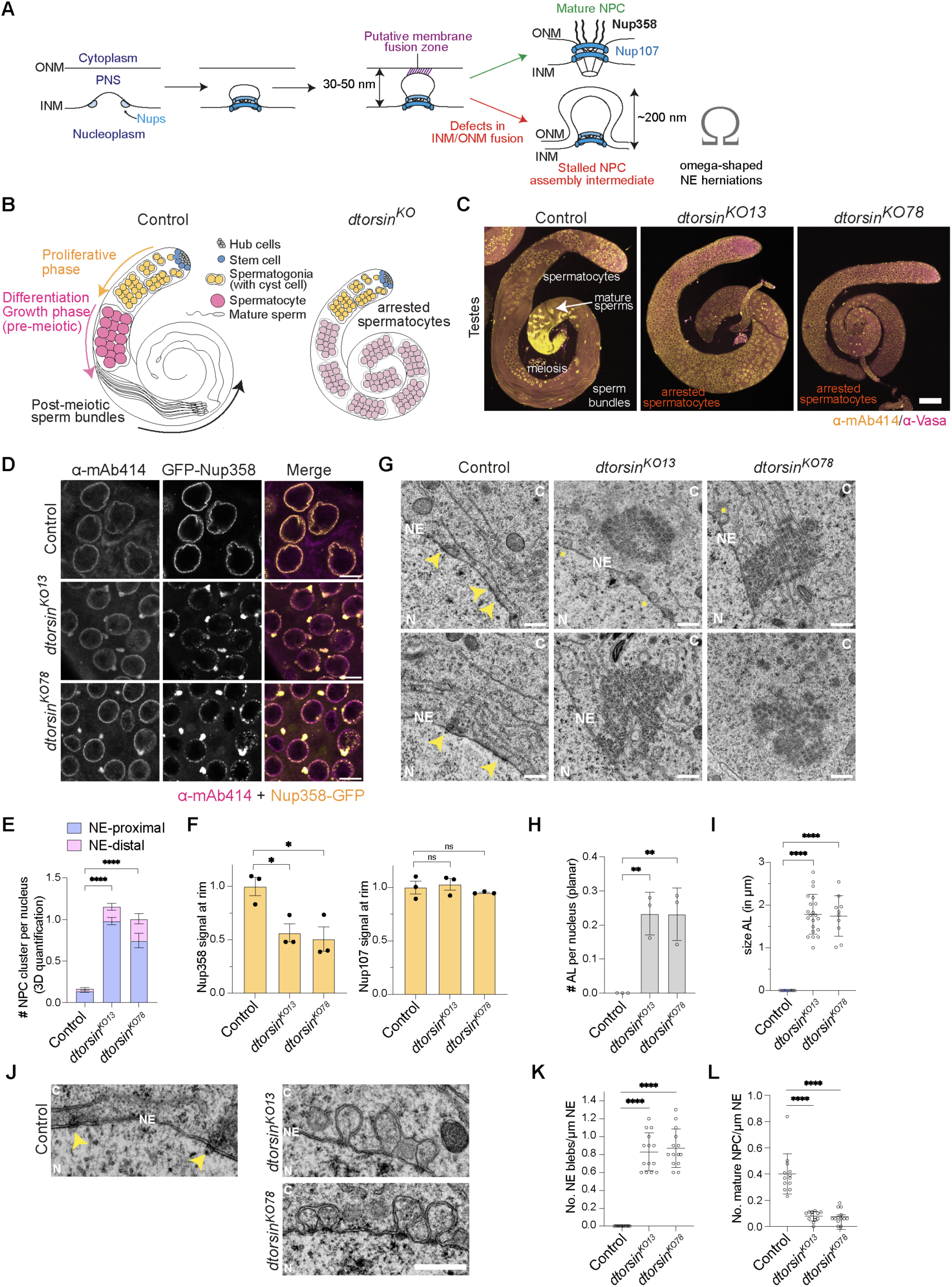
Spermatocytes of dtorsinKO flies are arrested in development, contain cytoplasmic NPC clusters reminiscent of annulate lamellae and display NPC biogenesis defects. **(A)** Schematic of NE remodelling and membrane fusion during interphase NPC biogenesis. Defects in membrane fusion, e.g. upon dysfunction of the Torsin system, manifest in NE herniations (‘blebs’) with a stalled NPC assembly intermediate at their constricted neck. Note that the cytoplasmic filaments formed by GFP-Nup358 are only added after fusion of the INM and ONM. **(B)** Scheme depicting the development of male germ cells in the *Drosophila* control and *dtorsin^KO^* germlines. **(C)** Representative confocal images of testes from control and two *dtorsin^KO^* strains (*dtorsin^KO13^*and *dtorsin^KO78^*), immunostained with mAb414 (anti-FG-Nups, yellow) and anti-Vasa (magenta) antibodies. Scale bar: 100 µm. **(D)** Representative confocal images of spermatocytes from either wild-type or two *dtorsin^KO^* strains expressing GFP-tagged Nup358 and immunostained with mAb414 (anti-FG-Nups) antibodies. N = 3. Scale bars: 10 µm. **(E)** Quantification of the number of FG-Nup-positive foci using three-dimensional (3D) reconstructed images (N = 3, n = 150, **** p ≤ 0.0001, student’s unpaired t-test, two-tailed, error bars, mean +/- SEM). **(F)** Quantification of the GFP-Nup358 signal from (C) and Nup107-GFP signal (from S2E) at the NE (N = 3, n = 60; one-way ANOVA, * p ≤ 0.05, mean +/-SEM). Data were normalized to the mean of the control group. **(G)** Representative images of annulate lamellae (AL)-like structures in *dtorsin^KO^* spermatocytes observed by transmission electron microscopy (TEM). Yellow arrowheads indicate mature NPCs. Yellow asterisks point to NE herniations present exclusively in *dtorsin^KO^* spermatocytes. C – Cytoplasm, N – Nucleus, NE – Nuclear Envelope. Scale bars: 500 nm. **(H)** Quantification of the number of AL per nucleus from (F). The analysis was performed on 2D slices taken 85 µm, 105 µm and 145 µm from the tip of the testis (n ≥ 50 cells, ** p ≤ 0.01, student’s unpaired t-test, two-tailed, mean +/- SD). **(I)** Quantification of the AL size from (F) (n ≥ 50 cells, **** p ≤ 0.0001, student’s unpaired t-test, two-tailed, mean +/- SD). **(J)** Representative TEM images of NE herniations at the nuclear envelope (NE) in *dtorsin^KO^* spermatocytes. Yellow arrowheads indicate mature nuclear pores in control spermatocytes. C – Cytoplasm, N – Nucleus. NE – Nuclear Envelope. Scale bar: 500 nm. **(K)** Quantification of the NE herniation number per µm of the NE from (J) (n = 13 cells, **** p ≤ 0.0001; student’s unpaired t-test, two-tailed; error bars, SD). **(L)** Quantification of the number of mature NPCs per µm of the NE from (J) (n = 13 cells, **** p ≤ 0.0001; student’s unpaired t-test, two-tailed; error bars, SD).

Torsin function is typically required for the viability and development of metazoans (Rose et al., 2015), and Tor1a−/− null mice die about 48 hours after birth (Goodchild et al., 2005). In contrast, ablation of the *Drosophila melanogaster* Tor1A homolog dTorsin is only ‘semi-lethal’ and about 10% of *dtorsin^KO^*flies escape pupal lethality (Wakabayashi-Ito et al., 2011). These escapers are males and develop to adulthood but are sterile, offering a unique experimental system to study the function of Torsins during male germ cell development.

Here, we demonstrate that the sterility of *dtorsin^KO^*males is explained by a developmental arrest of rapidly growing germ cells known as spermatocytes. Immature *dtorsin^KO^* spermatocytes show defects in the fusion of the INM and ONM, thereby compromising the final steps of NPC biogenesis at the NE. By exploiting proximity labelling and AlphaFold modelling, we identify the human ER membrane protein CLCC1 and its fly homolog CG12945 (named dClcc1) as novel Torsin interaction partners. Characterization of CLCC1 and dClcc1 reveals that their depletion phenocopies the INM-ONM fusion defects observed upon Torsin inactivation. The localization of CLCC1 at designated membrane fusion sites supports its direct role in the fusion process. In addition, mutations of dTorsin at the dClcc1 interaction interface fail to rescue the phenotypes of d*torsin^KO^* spermatocytes, showcasing the functional importance of this association. Remarkably, increased expression of CLCC1 or dClcc1 is sufficient to rescue the cellular defects of Torsin deficiency in human somatic or *Drosophila* germ cells, respectively. Taken together, our work identifies CLCC1 as a candidate for the long sought-after Torsin-regulated NE membrane fusogen, informing a mechanistic model of membrane fusion at the NE. Our findings also imply that tuning CLCC1 expression may be a promising therapeutic strategy for DYT1 dystonia.

## Results

### Arrested development of *dtorsin^KO^* germ cells is associated with NPC biogenesis defects at the NE and the formation of annulate lamellae

*D. melanogaster* harbours three genes encoding for Torsin-like proteins of which dTorsin/CG3024 is most closely related to human Tor1A (Fig. S1A). Deletion of *dtorsin* is semi-lethal and those male animals that do escape pupal lethality are sterile (Wakabayashi-Ito et al., 2011). To analyse the cause of male sterility associated with the loss of dTorsin, we stained testes from control and two independent *dtorsin^KO^* lines (*dtorsin^KO13^* and *dtorsin^KO78^* (Wakabayashi-Ito et al., 2011)) for Vasa (a germ cell marker) and FG-repeat rich nucleoporins (FG-Nups) of NPCs (using mAb414, which highlights the nuclear contours). In control testes, we observed germ cells across all stages of gametogenesis, including the proliferative mitotic stages (spermatogonia), the pre-meiotic stages (spermatocytes) and the mature sperm bundles at the distal tip of the testes (Fig. 1B, C). In contrast, we observed a failure in spermatocyte maturation in *dtorsin^KO^* testes, i.e., a striking developmental arrest that underlies the sterility of these mutant animals. These arrested spermatocytes, which normally grow ∼25-fold during germline development (Yamashita, 2018), also exhibited significantly reduced nuclear volumes in the absence of dTorsin function (Fig. S2A, B).

Since Torsins have been implicated in NPC biogenesis (Jacquemyn et al., 2021, Kim et al., 2024, Laudermilch et al., 2016, Pappas et al., 2018, Rampello et al., 2020) (Fig. 1A), we examined the distribution patterns of nucleoporins (Nups) in the arrested *dtorsin^KO^* spermatocytes, revealing major phenotypes. Firstly, immunostaining of bulk FG-Nups with mAb 414 (Fig. 1D, E, F) and analysis of the FG-Nup Nup58-GFP (Fig. S2C) both indicated that *dtorsin^KO^* spermatocytes contained on average one prominent perinuclear cluster of FG-Nups in the cytoplasm. The perinuclear FG-Nup foci in *dtorsin^KO^* spermatocytes were also positive for GFP-Nup107, a scaffold Nup located in the cytoplasmic and nuclear rings of NPCs (Fig. S2D, Movies S1, S2), indicating that these clusters do not merely represent FG-Nup aggregates. Notably, the Nup foci were also enriched in the RNA helicase Vasa (Fig. S2D), which is known to localize to germ cell RNA-rich granules at the nuclear periphery (Pamula & Lehmann, 2024). Similarly, the ovaries of homozygous *dtorsin^KO^* females that we generated were rudimentary, with egg chambers that did not develop past stages 4-5 (Fig. S3A, B, C), ultimately leading to female sterility (Fig. S3D, E). As in spermatocytes, perinuclear FG-Nup positive foci were also observed in the arrested nurse cells of *dtorsin^KO^*ovaries (Fig. S3F, G, H). Importantly, rescue experiments showed that re-expression of dTorsin using either a *dtorsin*-containing chromosomal duplication (*Rescue^Dup^*) or an inducible *dtorsin* transgene (*Rescue^gene^*) suppressed Nup foci formation and restored fertility (Fig. S2E-H, Fig. S3D-H).

Secondly, the analysis of GFP-tagged Nup358, a large Nup that constitutes the cytoplasmic filaments of metazoan NPCs (Bley et al., 2022, Zhu et al., 2022) (Fig. 1A), revealed NPC assembly defects at the NE. While GFP-Nup358 was present in the perinuclear Nup foci of arrested *dtorsin^KO^* spermatocytes, its signal intensity at the NE was strongly diminished, whereas the Nup107 signal at the NE was not majorly affected (Fig. 1D, F). Since the addition of the cytoplasmic NPC filaments only occurs after the fusion of the inner and outer nuclear membranes and exposure of late NPC assembly intermediates to the cytoplasm (Fig. 1A), the reduction in Nup358 levels at the NE is indicative of defects in membrane fusion required for the final steps of interphase NPC assembly (Otsuka et al., 2016).

We next performed transmission electron microscopy (TEM) on *dtorsin^KO^*spermatocytes to gain insights into the ultrastructure of both the perinuclear Nup foci and potential NE aberrations. TEM analysis revealed the presence of electron-dense foci, which contained dozens of clustered NPCs, in the vicinity of the NE (Fig. 1G). The NPCs contained in these clusters were regularly spaced and embedded in stacked membrane sheets termed annulate lamellae (Kessel, 1963, Kessel, 1992, Merriam, 1959, Wischnitzer, 1970), which were approximately 2 µm in diameter (Fig. 1G-I). In addition, we observed the accumulation of omega-shaped membrane herniations (NE ‘blebs’) with NPC-like structures within their constricted necks at the NE of *dtorsin^KO^* spermatocytes (Fig. 1A, J, K). These NE blebs highlight a failure in nuclear membrane fusion and are the hallmark of Torsin loss-of-function (Demircioglu et al., 2019, Goodchild et al., 2005, Jacquemyn et al., 2021, Kim et al., 2010, Laudermilch et al., 2016, Liang et al., 2014, Naismith et al., 2009, Rampello et al., 2020, Tanabe et al., 2016, VanGompel et al., 2015). Both tested *dtorsin^KO^* strains exhibited about 8 NE herniations per 10 µm of NE, accompanied by a significant reduction in the number of mature NPCs (Fig. 1J, L).

To assess whether the reduction in mature NPCs affected nuclear transport, we examined poly(A)-mRNA export by fluorescence in situ hybridization (FISH). In *dtorsin^KO^* spermatocytes, the nuclear-to-cytoplasmic ratio of the poly(A)-RNA FISH signal was slightly increased, indicating a mild mRNA export defect (Fig. S2I, J). Notably, we also observed a poly(A)-RNA FISH signal at the annulate lamellae of the *dtorsin^KO^* spermatocytes (Fig. S2I), consistent with the presence of the RNA helicase Vasa in the Nup foci (Fig. S2D).

Together, these findings indicate that the lack of dTorsin impairs the final steps of NE-associated NPC biogenesis, resulting in the accumulation of NE herniations. This defect in NPC assembly at the NE is best explained by a failure of the membrane fusion step that exposes newly assembled NPCs to the cytoplasm for the completion of their assembly. At the same time, NPCs can assemble, in principle, in the ER, likely in preexisting holes formed during remodelling of the ER membrane network (Zucker & Kozlov, 2022) without the need for membrane fusion, giving rise to annulate lamellae.

### A conserved N-terminal cysteine cluster is required for dTorsin function at the Nuclear Envelope

To investigate which of dTorsin’s conserved features are required for its function, we performed rescue experiments with GFP-tagged dTorsin variants expressed in *dtorsin^KO^* germ cells, and examined their respective effects on annulate lamellae formation, sperm production and fertility. Surprisingly, constructs carrying mutations in the Walker A (K114A) and Walker B motifs (E177Q) of dTorsin partially rescued the *dtorsin^KO^* phenotypes (Fig. S4A-D), indicating that neither ATP binding nor hydrolysis by dTorsin are essential for germ cell development. We also deleted the entire gene encoding for the single fly Torsin ATPase activating protein TorIP (Torsin Interacting Protein) (Demircioglu et al., 2016, Jacquemyn et al., 2021, Sosa et al., 2014), which is the homolog of human LAP1 and LULL1 (Fig. S1A). Consistent with the above experiments, *torip^KO^* males were viable, fertile and lacked the prominent cytoplasmic Nup clusters in their spermatocytes (Fig. S4E-G). This confirms that Torsin ATPase activity is largely dispensable for germline development, extending previous observations in the fly fat body (Jacquemyn et al., 2021).

Torsins harbour other conserved features, including a conserved glycine residue (G256) at a potential interaction or oligomerization interface (Chase et al., 2017), and six conserved cysteine residues (Fig. S1B, C, Fig. S4A, H). Torsin carrying the G256D mutation was unable to rescue the phenotypes observed in the *dtorsin^KO^* testes (Fig. S4B-D), suggesting that higher order complex formation of dTorsin is critical for its core function. Similarly, dTorsin variants carrying combinatorial C-to-S mutations of the N-terminal cysteine cluster failed to suppress Nup foci formation and to rescue fertility, revealing their requirement for Torsin function (Fig. S4J-K). These N-terminal cysteines lie adjacent to a hydrophobic region that attaches the ATPase to the membrane and could be involved in forming disulfides, perhaps to correctly orient the luminal domain of Torsins for oligomerization or the interaction with other partners (Fig. S1C).

### The disease causing Torsin1AΔE mutant localizes to NE blebs at sites of membrane apposition

Interphase NPC biogenesis begins with the remodelling of the INM in conjunction with the assembly of the NPC scaffold (Otsuka et al., 2016), which generates the membrane pores in which NPCs are housed (Fig. 1A). Subsequently, membrane fusion of the INM and ONM occurs at the emerging bulge of the INM, which is positioned over the assembling NPC, to expose pre-NPCs to the cytoplasm for their final assembly steps (Otsuka et al., 2016). Considering the omega-shaped NE herniations that form upon Torsin inactivation, we wondered whether Torsins localise to the putative membrane fusion zone, i.e., the region of INM and ONM apposition that forms at the tip of NE herniations. To induce NE herniations and concomitantly examine Torsin localization, we expressed the dystonia-causing human Torsin1AΔE mutant as a GFP fusion in cells lacking LULL1 and LAP1 (*TOR1AIP1/TOR1AIP2* DKO), the absence of which is known to induce NE herniations (Laudermilch et al., 2016), and used a gold-coupled anti-GFP antibody for Torsin1AΔE-GFP detection by TEM. Torsin1AΔE-GFP was indeed highly enriched at the NE herniations in the perinuclear space, immediately next to the zone of INM-ONM apposition (Fig. 2A, B). This localisation strongly supports a direct role for Torsins in the membrane fusion reaction that is needed for both the final steps of NPC assembly and other nuclear egress pathways to the cytoplasm.

**Figure 2:**
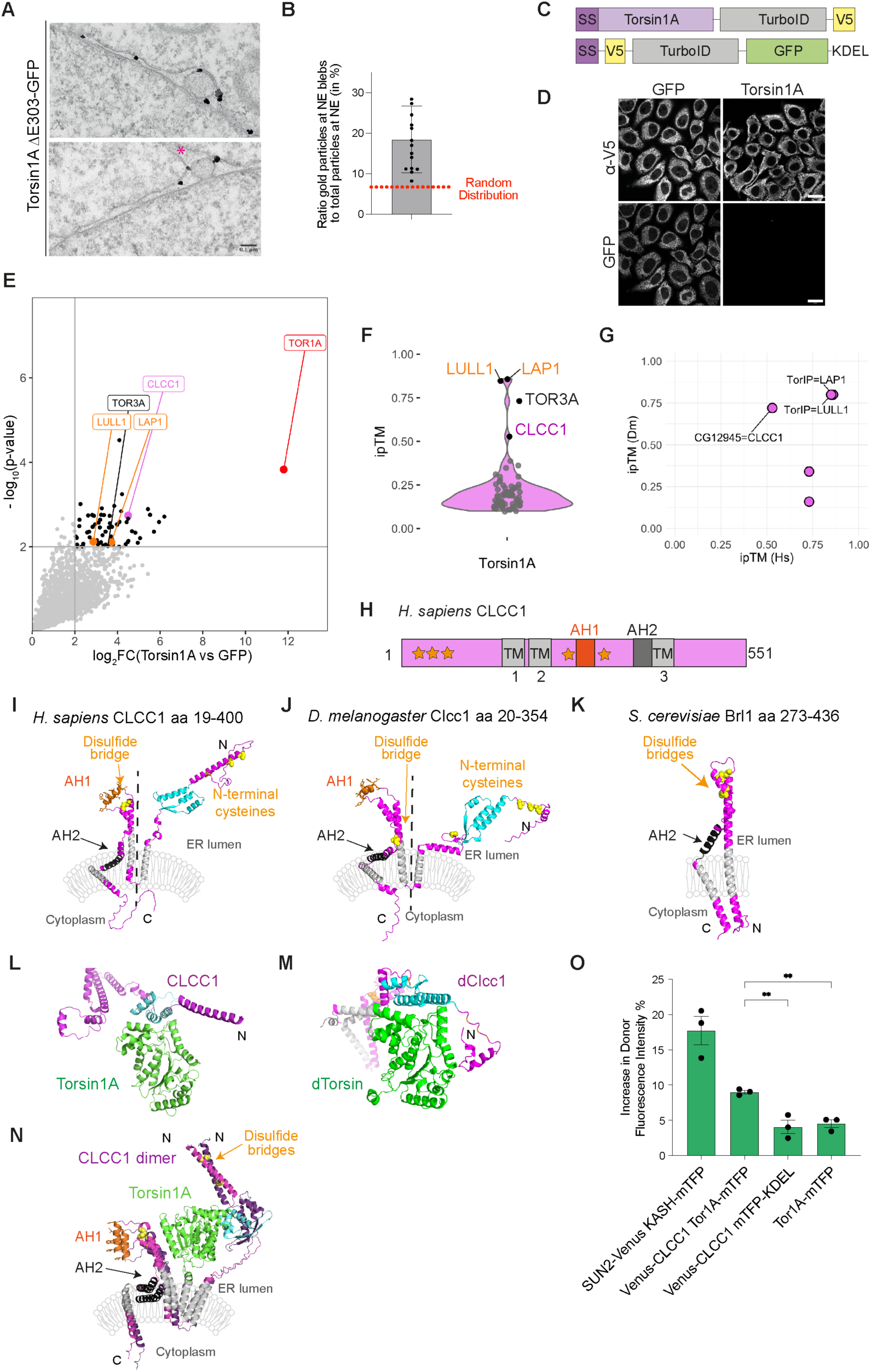
Proximity labelling of Torsin1A in HeLa cells identifies CLCC1 as a candidate direct interaction partner. **(A)** Immunogold localization of Torsin1AΔE302/303-GFP by TEM in gene-edited HeLa *TOR1AIP1/TOR1AIP2* KO cells (genes encoding for LAP1 and LULL1) using a gold-labelled anti-GFP antibody. Note the enrichment of gold particles at the sites of INM-ONM apposition within the NE blebs. Pink asterisks marks occasionally observed dome-like membrane structures above NE blebs. **(B)** Quantification of the fraction of gold particles enriched at NE blebs (herniations) relative to the total NE perimeter including NE herniations (n = 15 cells; mean +/- SD). The expected random distribution was estimated based taking into account bleb occupancy at the NE, relative to the total measured NE perimeter. **(C)** Scheme of the V5 epitope-tagged Torsin1A-TurboID and TurboID-GFP fusion proteins. TurboID-GFP (negative control) contained an ER signal sequence and a C-terminal KDEL sequence to ensure ER localization. **(D)** Representative confocal images of HeLa cells expressing V5-tagged Torsin1A-TurboID or ER-localized TurboID-GFP. Cells were immunostained using anti-V5 antibodies. Scale bars, 20 µm. **(E)** One-sided volcano plot of the ratio of biotinylated proteins identified on Torsin1A over GFP (N = 3). Proteins with adjusted p-values < 0.01 and log_2_(fold change, FC) > 2 (grey lines, ANOVA test and Benjamini–Hochberg procedure) were considered significant and are marked as black points. **(F)** Violin plot of ipTM scores obtained from AlphaFold-Multimer predictions of Torsin1A in complex with significant interactors identified in (E) with adjusted p-values < 0.01 and log_2_(fold changes) > 2. Black dots represent proteins with ipTM score > 0.5. **(G)** Correlation of ipTM scores of identified human proteins with ipTM > 0.5 (ipTM (Hs)) with the corresponding ipTM scores of their *Drosophila melanogaster* homologs (ipTM (DM)), obtained by HHpred. **(H)** Scheme of the human CLCC1 protein with depicted structural features. AH – amphipathic helix, TM – transmembrane domain. Asterisks mark cysteine residues. **(I, J, K)** Membrane topology models of the indicated parts of human CLCC1, *Drosophila* Clcc1 (CG12945) and *S. cerevisiae* Brl1 based on their AlphaFold3 predictions. A three-helical domain of CLCC1 and dClcc1, predicted to be involved in Torsin interaction, is highlighted in cyan. TM domains are shown in grey. Yellow dots represent cysteines and disulfide bridges. Transmembrane helices are in grey, AH – amphipathic helix. The dashed line separates the N-terminal Torsin binding region and the part of CLCC1/dClcc1 related to Brl1. **(L, M)** AlphaFold Multimer models of complexes between human Torsin1A (aa 40 - 334) and the N-terminal region of CLCC1 (aa 60 - 305) as well as of fly dTorsin (aa 40 - 340) and the N-terminal region of dCLCC1 (aa 20 - 330). CLCC1/Clcc1 shown in pink/cyan (as in I, J), Torsin1A and dTorsin in green. **(N)** AlphaFold Multimer model of human Torsin1A interacting with a CLCC1 dimer. AH1 is in orange, cysteines in yellow. **(O)** FRET efficiency between CLCC1 and Torsin1A determined by acceptor photobleaching after expression of the indicated fusion proteins in HeLa *CLCC1* KO cells. mTFP served as the donor fluorophore, Venus as the acceptor fluorophore. SUN2-Venus and mini-Nesprin2-mTFP (mTFP inserted between the TM segment and the KASH peptide) served as positive control (Frischer, 2019, Sosa et al., 2012). N = 3, n = 40 cells total per condition, one-way ANOVA, ** p ≤ 0.01, mean +/- SEM).

### Proximity labelling identifies CLCC1 as an evolutionary conserved interaction partner of Torsins

To shed light on the molecular function of Torsins in membrane fusion, we set out to identify Torsin binding partners. Traditional affinity purification methods have identified LAP1 and LULL1 as major Torsin interactors (Goodchild & Dauer, 2005). However, potential Torsin substrates, including the enigmatic INM-ONM fusogen, have eluded discovery, likely due to the loss of binding partners during the purification procedure. To overcome this limitation, we employed TurboID-based biotin proximity labelling (Branon et al., 2018) in HeLa cell lines expressing Torsin1A fused to the biotin ligase TurboID, or an ER-luminal TurboID-GFP as a negative control (Fig. 2C, D). Mass spectrometry analysis identified approximately 70 proteins that were enriched on Torsin1A compared to the GFP control (Fig. 2E; Table S1; adj. p-value < 0.01, log_2_FC > 2,). The Torsin activators LAP1 and LULL1 were among the interactors, confirming the validity of the method.

To identify potential direct interactors among the ∼70 top candidates, we applied AlphaFold Multimer (AFM) modelling and determined the interphase predicted Template Modelling scores (ipTM), which provide a numerical measure of confidence for the predicted interfaces (Fig. 2F) (Evans et al., 2022). As expected, LAP1 and LULL1 displayed ipTM scores higher than 0.75 (good confidence). Among the few other candidates with moderate confidence ipTMs scores (> 0.5) was the Chloride Channel CLIC-like protein 1 (CLCC1) (Fig. 2F). Reassuringly, we also found a CLCC1- Torsin1A/B complex to be predicted with very high confidence by a recent deep learning approach applied to the human proteome ((Zhang et al., 2025), (Zhang et al., 2024). In support of a direct interaction between CLCC1 and Torsin1A, we observed Förster Resonance Energy Transfer (FRET) between Venus-CLCC1 and Tor1A- mTFP expressed in HeLa cells (Fig. 2O), but slightly less efficient than for our positive control, a SUN-KASH FRET pair, a stable, disulfide-linked complex that builds the centrepiece of LINC complexes (Frischer, 2019, Sosa et al., 2012).

Although CLCC1 was first described as an ion channel protein (Guo et al., 2023), it has recently been linked to cellular lipid transport (Mathiowetz et al., 2024, Wu et al., 2024). Its central domain possesses structural similarity to the yeast proteins Brl1 and Brr6 (Fig. 2I, K) (Mathiowetz et al., 2024), both also implicated in lipid metabolism and NPC biogenesis (Kralt et al., 2022, Lone et al., 2015, Vitale et al., 2022). Interestingly, CLCC1 homologs have also been discovered in the genome of herpes viruses and suggested to promote viral nuclear egress (Dai et al., 2024), a process that requires INM-ONM fusion.

If CLCC1 is a key interaction partner of Torsins across metazoan evolution, then CLCC1 should also exist in flies. By exploiting the structure alignment tool HHpred, we identified an uncharacterized protein encoded by the CG12945 gene as the putative *Drosophila* CLCC1 homolog (hereafter dubbed dClcc1). The dTorsin-dClcc1 complex obtained an AFM ipTM score of ∼ 0.75, similar to established complexes of human Torsin1A with LAP1 and LULL1 or the predicted dTorsin-TorIP complex (Fig. 2G). Furthermore, Hidden Markov Model-based homology searches across eukaryotic proteomes revealed that CLCC1 and Torsin homologs co-occur throughout the metazoan lineage, supporting the model of a functional association conserved across 600 million years of animal evolution (Fig. S5A, B).

CLCC1 and dClcc1 are strikingly similar at the structural level (Fig. 2H-J), despite sharing only ∼ 22% sequence identity. Both proteins feature an N-terminal luminal domain predicted to be involved in Torsin interaction (Fig. 2I, J, L, M), which is lacking in the yeast Brl1-like proteins (Fig. 2K, (Fischer et al., 2025)), consistent with the absence of Torsin in yeasts (Fig. S5). The N-terminal regions of both CLCC1 and dClcc1 contain cysteines, although their exact position seems poorly conserved (Fig. S5D). These cysteines may promote CLCC1 dimerization by the formation of interchain disulfide bonds (Fig. S6A) (Guo et al., 2023, Mathiowetz et al., 2024). AF modelling also predicts that CLCC1 can form higher order, ring-shaped assemblies of 10 or more protomers (Fig. S6C, D) (Dai et al., 2024, Guo et al., 2023, Mathiowetz et al., 2024, Wu et al., 2024), akin to what has been suggested for Brl1 and Brr6 (Fischer et al., 2025). Both the dimeric (Fig. 2N) and higher order multimers of CLCC1 (Fig. S6E) are structurally compatible with Torsin interaction.

#### Loss of dClcc1 phenocopies dtorsin^KO^ loss-of-function revealing its role in NE membrane fusion during NPC biogenesis

To test whether Torsins and CLCC1 are involved in the same pathway, we examined whether loss of dClcc1 phenocopies the *dtorsin^KO^* phenotypes. First, we generated a *dClcc1* knockout allele (*dclcc1^KO^*) and found that homozygous *dclcc1^KO^* animals could not be obtained, indicating that dClcc1 is essential for the viability of flies. To circumvent organismal essentiality, we used the FLP-FRT system to generate mosaic animals whose testes contain a mixture of dClcc1-containing and *dclcc1^KO^* germ cell cysts that can be distinguished based on the differential expression of an RFP-NLS fusion protein (Fig. 3A). Akin to dTorsin loss-of-function, we observed that *dclcc1^KO^* spermatocytes (RFP-negative), but not their dClcc1-containing neighbours (RFP-positive), exhibited perinuclear foci that also contained Vasa (Fig. 3B, C). Similar perinuclear Nup foci were also found in spermatocytes homozygous for a P-element insertion allele of *dclcc1* (*dclcc1^EP^*) (Fig. 3D, E). In addition, these *dclcc1^EP^*spermatocytes exhibited reduced GFP-Nup358 fluorescence at the nuclear rim (Fig. 3D, E), indicative of a defect in the completion of NPC assembly. At the same time, EGFP-Nup358 was present in NPCs of the juxtanuclear annulate lamellae, akin to what was observed in *dtorsin^KO^* spermatocytes (Fig. 1D).

**Figure 3:**
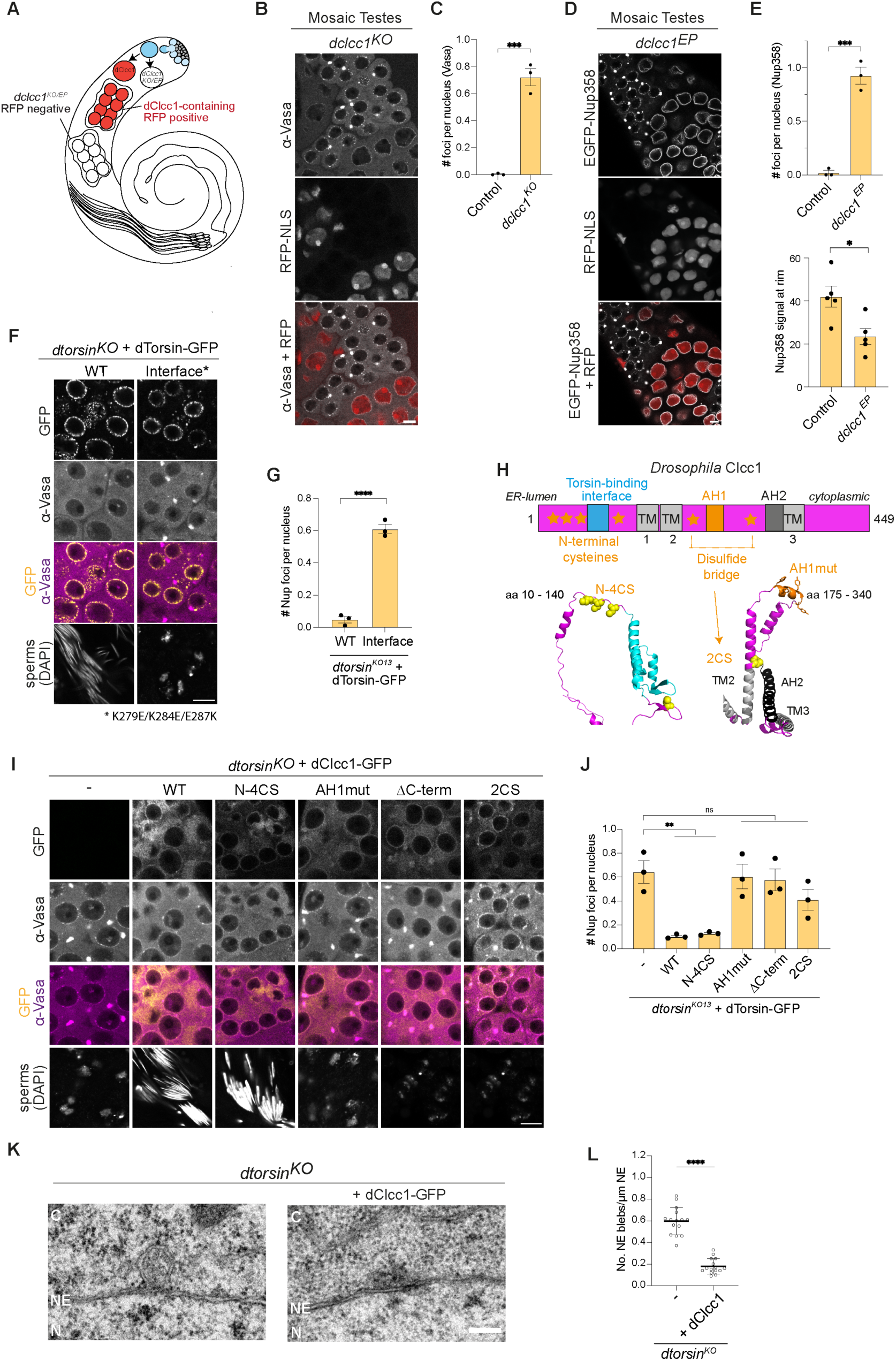
Loss of *Drosophila dClcc1* phenocopies the NPC biogenesis defects caused by *dtorsin* loss-of-function. **(A)** Schematic of a testis tissue mosaic with dClcc1-containing (RFP-positive) and *dClcc1^KO^*/*dClcc1^EP^*(RFP-negative) spermatocytes generated using the FLP-FRT system. RFP-positive cells contain dClcc1 while the RFP-negative cells are homozygous for the *dclcc1^KO^* or *dclcc1^EP^* alleles. *dclcc1^EP^* flies constitutively express EGFP-Nup358. **(B)** dCLCC1-deficient spermatocytes form Vasa-positive foci. Mosaic *dclcc1^KO^* testes (RFP-positive cells contain dClcc1, RFP-negative cells are homozygous for the *dclcc1^KO^*allele) were immunostained with an anti-Vasa antibodies. Scale bar: 10 µm. **(C)** Quantification of the number of Vasa containing foci in RFP-positive (control) and RFP-negative cells in (B) (N = 3, n ≥ 123 cells, *** p ≤ 0.001, student’s unpaired t-test, two-tailed, mean +/- SEM). **(D)** dCLCC1-deficient spermatocytes in the mosaic testes of a *dclcc1^EP^* line (RFP-positive cells contain dClcc1 while the RFP-negative cells are homozygous for the *dclcc1^EP^* allele) constitutively expressing EGFP-Nup358 form Nup358-positive annulate lamellae and display defects in final steps of NPC biogenesis. Scale bar: 10 µm. **(E)** Quantification of the number of EGFP-Nup358 containing AL (top panel: N = 3; n ≥ 123 cells; *** p ≤ 0.001; student’s unpaired t-test, two-tailed, mean +/- SEM) and of the EGFP-Nup358 signal at the NE (bottom panel: N = 5, n ≥ 70 cells, * p ≤ 0.05, student’s unpaired t-test, two-tailed, mean +/- SEM) (mean fluorescence intensity values, arbitrary units) from (D) in RFP-positive (control) and RFP-negative *dclcc1^EP^*spermatocytes. **(F)** A dTorsin mutant carrying mutations (K279E/K284E/E287K, interface mutant) at the dClcc1 binding interface does not rescue defects in *dtorsin^KO13^* spermatocytes, whereas wildtype dTorsin does. dTorsin-GFP variants were expressed under the control of *bam-Gal4* in *dtorsin^KO13^*testes, which were stained for Vasa (magenta) and DAPI (white). Zoom representation of the dTorsin and dClcc1 interaction interface with key interacting residues. Scale bar: 10 µm. **(G)** Quantification of the number of Vasa-containing AL from (E) (N = 3, n ≥ 270 cells, **** p ≤ 0.0001, student’s unpaired t-test, two-tailed, mean +/- SEM). **(H)** A schematic of *Drosophila* Clcc1 (dClcc1) protein with the depicted structural features. AH – amphipathic helix, TM – transmembrane domain. Mutated features include the N-terminal cysteines (mutant: N-4CS), the disulfide bridge (2CS) and the aromatic residues in amphipathic helix 1 (AH1mut). **(I)** Expression of the indicated *Drosophila* dClcc1-GFP variants (yellow) under the control of *bam-Gal4* in *dtorsin^KO13^* testes stained for Vasa (magenta) and DAPI (white). WT – wildtype dClcc1, N-4CS – cysteine-to-serine mutations in four N-terminal cysteines (C48S, C50S, C53S, C140S), AH1mut – mutations in amphipathic helix 1 (W234S/F235S/Y241S/Y243S), ΔC – deletion of the cytoplasmic (C-terminal) region (Δ341-449), 2CS – cysteine-to-serine mutations of two conserved cysteines on both sides of the AH1 (C211 and C266) forming a disulfide bridge. **(J)** Quantification of the number of Vasa-containing AL from (H) (N = 3, n ≥ 198 cells, ** p ≤ 0.01, ns p > 0.05, student’s unpaired t-test, two-tailed, mean +/- SEM). **(K)** TEM analysis of *dtorsin^KO13^*spermatocytes without and with expression of additional dClcc1-GFP. C – Cytoplasm, N – Nucleus, NE – Nuclear Envelope. Scale bar: 200 nm. **(L)** Quantification of the number of NE herniations in *dtorsin^KO13^* spermatocytes without and with expression of additional dClcc1-GFP from (J) (n ≥ 15, **** p ≤ 0.0001, student’s unpaired t-test, two-tailed, mean +/- SD).

Having established that both Torsin and CLCC1 are required for the final steps of NPC biogenesis at the NE, we wished to further explore their functional relationship. The predicted Torsin binding interface of CLCC1/dClcc1 involves a three-helical bundle (Fig. 2L, M; cyan), which engages with the folded Torsin ATPase domain through salt bridges (Fig. S6F). To examine whether the function of dTorsin depends on its interaction with dClcc1, we mutated three residues in dTorsin (K279E/K284E/E287K) in the predicted binding interface (Fig. 3F). Strikingly, the dTorsin interface mutant failed to suppress annulate lamellae formation and to rescue spermatogenesis, despite being expressed and targeted to the NE like wildtype dTorsin (Fig. 3F, G). Together, these results support the notion that a direct physical interaction between dClcc1 and dTorsin is required for NE membrane fusion during NPC biogenesis.

### Increased levels of dClcc1/CLCC1 suppress Torsin loss-of-function phenotypes

To better understand how Torsin and CLCC1 function together, we expressed a dClcc1-GFP fusion protein on top of endogenous dClcc1 in dTorsin-lacking germ cells. Remarkably, increased dClcc1 expression was sufficient to reduce annulate lamellae formation in *dtorsin*^KO^ spermatocytes and restore spermatogenesis in these animals (Fig. 3H-J, compare (-) vs. WT dClcc1). Furthermore, ultrastructural TEM analysis demonstrated that the formation of NE herniations was also suppressed by increased dClcc1 expression (Fig. 3K, L).

Similarly, we tested whether increased expression of human CLCC1 was able to suppress the accumulation of NE herniations in Tor4KO HeLa cells lacking four Torsin family members (Tor1A, Tor1B, Tor2A, Tor3A; (Rampello et al., 2020)). In these cells, NE blebs caused by a failure in INM-ONM fusion can be conveniently identified by immunostaining of either K48-linked ubiquitin or the J-protein cofactor MLF2 (Rampello et al., 2020). While ubiquitin-positive puncta at the NE were prominent in the Tor4KO cells (Fig. 4A), expression of CLCC1-GFP abolished the NE blebs (Fig. 4A, B), in agreement with the fly data. Collectively, these data demonstrate that excess dClcc1/CLCC1 makes the presence of Torsin(s) dispensable. Importantly, this suggests that the main function of Torsins is to sustain CLCC1 functionality.

**Figure 4:**
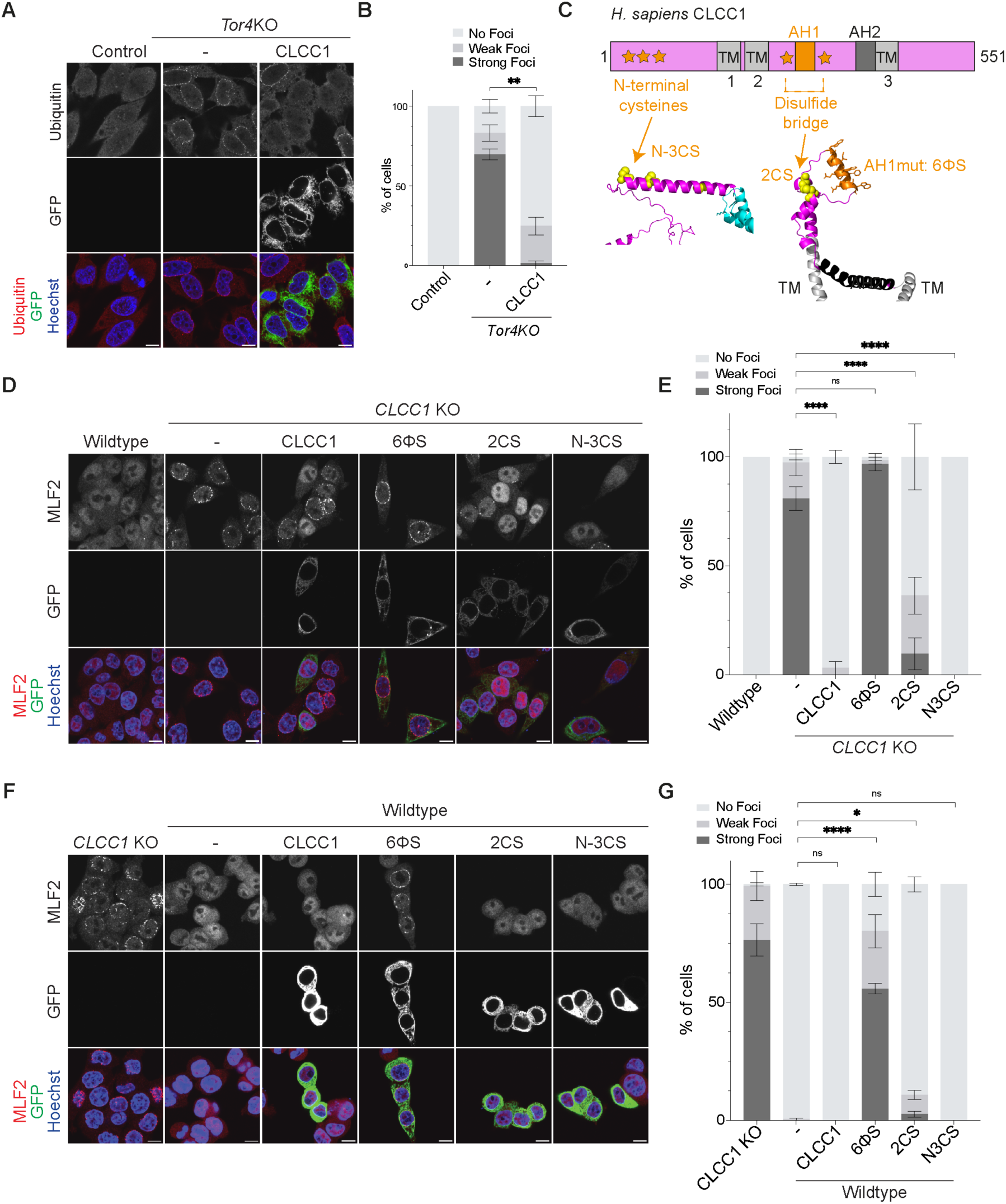
CLCC1 expression rescues defective NPC biogenesis in human Torsin knockout cells and identification of key functional elements in CLCC1. **(A)** Increased expression of CLCC1 in Tor4KO cells suppresses ubiquitin accumulation at the NE. Representative confocal images of Tor4KO HeLa cells that were transiently transfected with vectors for expression of CLCC1-GFP, fixed 48 h after transfection, immunostained for ubiquitin, stained with Hoechst and analysed by confocal microscopy. Scale bar: 10 μm. **(B)** Quantification of ubiquitin foci in (A), determining the fraction of control, Tor4KO, and Tor4KO expressing CLCC1-GFP cells with strong, weak or no ubiquitin foci (N = 3, n ≥ 80, ** p ≤ 0.01, student’s unpaired t-test, two-tailed, mean +/- SEM). **(C)** Scheme of the human CLCC1 protein illustrating the location of key structural features (AH – amphipathic helix, TM – transmembrane domain, asterisks – cysteines) plus AlphaFold model of the N-terminal CLCC1 domain indicating features that were targeted by mutational analysis. AH1 mutant 6<1S: W260S, I264S, W265S, W267S, F268S, W272S; disulfide bridge mutant: 2CS; N-terminal cysteine mutants: N-3CS. **(D)** Representative confocal images of suppression of MLF2 foci formation in HCT116 *CLCC1* KO cells by expression of wildtype CLCC1-GFP wildtype, disulfide and N-terminal cysteine mutants but not by a mutant in AH1 (CLCC1^6ΦS^-GFP). HCT116 CLCC1 KO cells were transfected with the indicated vectors, fixed 48 h after transfection, immunostained with anti-MLF2 antibodies, and analysed by confocal microscopy (N = 3). Scale bar: 10 μm. **(E)** Quantification of experiment in (D), determining the fraction of WT, *CLCC1* KO cells, and CLCC1-GFP re-expressing KO cells showing strong, weak or no MLF2 foci (N = 3; n ≥ 60; one-way ANOVA; **** p ≤ 0.0001). **(F)** Representative confocal images of HCT116 *CLCC1* KO and WT HCT116 cells as well as WT HCT116 cells expressing the indicated CLCC1-GFP variants. Cells were transfected with the indicated vectors, fixed 48 h after transfection, immunostained with anti-MLF2 antibodies, and analysed by confocal microscopy (N = 3). Note that the AH1 mutant CLCC1^6ΦS^-GFP exhibits a dominant-negative effect and induces MLF2 foci formation. Scale bar: 10 μm. **(G)** Quantification of experiment in (F), determining the fraction of *CLCC1* KO, wildtype and CLCC1-GFP re-expressing WT cells showing strong, weak or no MLF2 foci (N = 3; n ≥ 60; one-way ANOVA; **** p ≤ 0.0001).

### Conserved molecular features promote CLCC1/dClcc1 function

A common hallmark of the CLCC1 and Brl1 protein families is an ER-luminal loop region that is organized by one or two disulfide bridges (Fig. 2I-K). In both cases, the loop contains a stretch of hydrophobic residues (positioned between TM2 and TM3 of CLCC1, and TM1 and TM2 of Brl1). While in CLCC1-like proteins the respective hydrophobic residues are mostly aromatic and part of a short amphipathic helix (AH1, Fig. 2I, K; Fig. S5C, they are aliphatic in Brl1-type proteins in fungi (Fischer et al., 2025). To examine the importance of the conserved loop region and other key structural features of CLCC1/dClcc1, we expressed select mutant transgenes in *dtorsin^KO^* germ cells and human *CLCC1* KO cells, respectively. In *Drosophila* (Fig. 3H-J), we found that dClcc1-GFP transgenes carrying mutations in AH1 (dClcc1^AH1mut^- GFP, W234S/F235S/Y241S/Y243S) or its organizing disulfide bridge (dClcc1^2CS^-GFP) failed to rescue the *dtorsin^KO^* phenotypes, indicating that the loop region containing the short amphipathic helix AH1 is essential for the function of dClcc1. We also observed that a dClcc1 mutant lacking the C-terminal domain (dClcc1^ΔC^-GFP) did not rescue the *dtorsin^KO^* phenotypes, showing an important contribution of the cytoplasmic domain (residues 341-449 following TM3). Interestingly, mutation of the N-terminal cysteines (dClcc1^4CS^-GFP), which potentially promote dClcc1 dimerization, are not required for rescue, consistent with the rapid evolution of the N-terminal domain of CLCC1.

To extend our functional analyses to human cells, we used human HCT116 *CLCC1* KO cells that we have recently generated, in which we observed NPC biogenesis defects and omega-shaped NE herniations (Fischer et al., 2025). Similar to Tor4KO cells (Rampello et al., 2020), HCT116 *CLCC1* KO cells showed a striking accumulation of MLF2 foci at the nuclear rim (Fig. 4C-E). The MLF2 foci phenotype could be efficiently suppressed by the expression of a wildtype CLCC1-GFP fusion protein. In contrast, a mutant CLCC1-GFP construct carrying substitutions in AH1 (AH1 mutant 6<1S: W260S, I264S, W265S, W267S, F268S, W272S) did not rescue the *CLCC1* KO phenotype (Fig. 4D, E), but even exhibited a dominant-negative phenotype when expressed in HCT116 wildtype cells (Fig. 4F, G). Mutation of the disulfide bridge at the base of the protein loop that contains AH1 (CLCC1^2CS^-GFP) had only minor consequences and largely, although incompletely, rescued the MLF2 foci (Fig. 4D, E). Similar to our observations in the fly system, mutation of cysteines within the N-terminal luminal region in the region CLCC1^N-3CS^-GFP did not impair rescue. Collectively, our mutational analyses revealed an evolutionarily conserved key contribution of the AH1-containing loop region to the function of CLCC1 proteins in NPC biogenesis.

To understand the potential role of the AH1 containing loop for CLCC1 function, we next performed molecular dynamics (MD) simulations of the human CLCC1 16-mer oligomer embedded in a model membrane bilayer. We tested two potential models that are compatible with the experimental data: the first one in which only one CLCC1 oligomer is embedded in the ONM, facing a curved INM evagination above an NPC assembly intermediate (Fig. 5A-C); the second one in which two distinct CLCC1 16- mer complexes reside in both INM and ONM and approach each other (Fig. S8A, B). In both cases, we observed that the presence of CLCC1 leads to tethering of the two membranes, mediated either by direct interactions of the AH1 with the opposite membrane in the model with only one oligomer (Fig. 5A) or by AH1-AH1 interactions in the model with two CLCC1 oligomers (Fig. S8A). In both instances, this tethering is associated with significant lipid remodelling and destabilization of the bilayer structure within the CLCC1 central pore, similar to simulations of the yeast orthologues Brl1 and Brr6 (Fischer et al., 2025). Overall, our *in silico* data support our mechanistic hypothesis that CLCC1 can work as a membrane tethering fusogen between the INM and the ONM via the activity of its AH1 helix.

**Figure 5:**
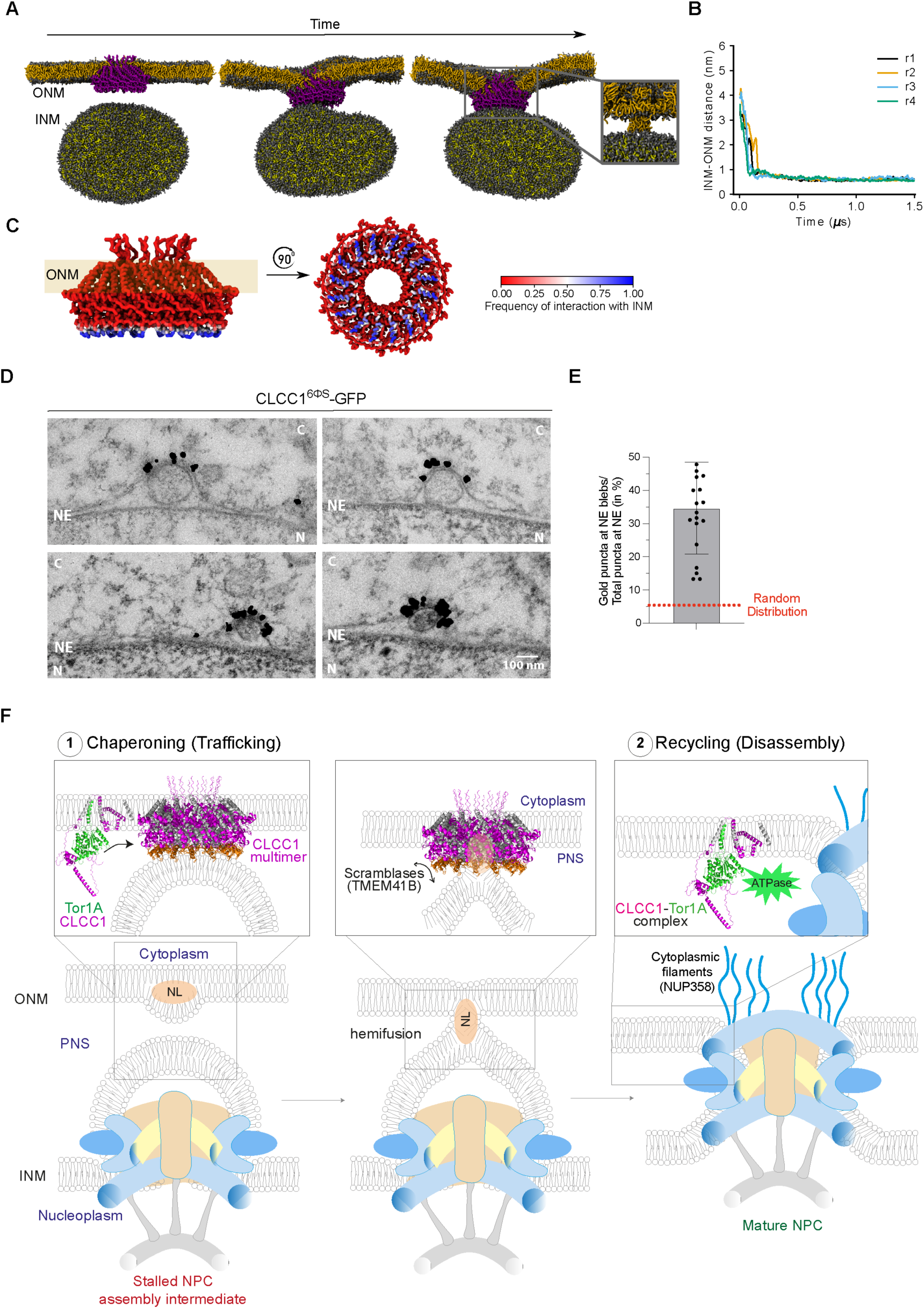
Mechanistic model of CLCC1 regulation by Torsin. **(A)** Coarse-grained MD simulation snapshots of a planar bilayer with CLCC1 16-mer inserted (ONM) and in opposition to a lipid vesicle (INM). Inset in the right most image shows ONM lipids in contact with INM vesicle but unable to mix with the lipids there. **(B)** Time-trace of minimum INM-ONM distance during the simulation showing stable contact between lipids of opposing membranes in (A). **(C)** CLCC1 with residues coloured according to the interaction frequency with the opposing membrane (INM) in side view (left) and top view (right) showing very high probability of AH1 helices to interact and insert into the opposing membrane. **(D)** Immunogold localization of a GFP-tagged CLCC1 AH1 mutant (CLCC1^6ΦS^-GFP) expressed in *CLCC1* KO HeLa cells using a gold-labelled anti-GFP antibody by TEM. The GFP epitope on the CLCC1 C-terminus resides in the cytoplasm. Note the strong accumulation of gold particles at the ONM-face of presumptive membrane fusion sites of NE blebs. **(E** Quantification of the fraction gold puncta enriched at the NE blebs (herniations) relative to the total NE perimeter including NE herniations. The expected random distribution (5.1%) was estimated based on the bleb occupancy at the NE relative to the total measured NE length (n = 20 cells, mean +/- SD). **(F)** Model of CLCC1-mediated ONM and INM fusion and CLCC1 regulation by Torsin. CLCC1 enriches at the designated fusion site at membrane blebs over NPC assembly intermediates. It forms oligomeric rings that house a central pore for lipid accumulation and exchange, potentially supporting the accumulation of neutral lipids (e.g. TAGs) between membrane leaflets to drive initial membrane deformation of the ONM, likely supported by other factors. This deformation may allow the close apposition of membranes to induce hemi-fusion. We envision two scenarios for Torsin regulation: (1) **Chaperoning/trafficking model**: CLCC1 traffics to sites of stalled NPC intermediates in complex with Torsin, which promotes proper multimer assembly at fusion sites. (2) **Recycling model**: Torsins disassemble CLCC1 oligomers, allowing recycling of subunits. Dismantling of CLCC1 oligomers may be required for the completion of membrane fusion. Gray helices mark transmembrane regions. NL – accumulation of neutral lipids; NPC – nuclear pore complex; INM – inner nuclear membrane; ONM – outer nuclear membrane; PNS – perinuclear space.

### The CLCC1 AH1 mutant accumulates at sites of INM-ONM apposition at NE blebs

CLCC1 has recently been implicated in lipid-related processes such as bilayer equilibration and hepatic neutral lipid flux (Mathiowetz et al., 2024, Wu et al., 2024). Since our data strongly suggests a direct function of CLCC1 in membrane fusion at the NE, we wondered whether it indeed localizes to NE herniations across NPC assembly intermediates. To test this hypothesis, we exploited the AH1 mutant (CLCC1^6ΦS^-GFP) to potentially trap CLCC1 at INM-ONM fusion sites. To examine whether it indeed accumulates at the tip of NE blebs in *CLCC1* KO cells, we employed immunogold microscopy to detect its C-terminal, extraluminal GFP moiety. Strikingly, CLCC1^6ΦS^-GFP was highly enriched at the cytoplasmic face of the NE blebs, with some herniations showing a prominent accumulation of gold particles at putative INM-ONM fusion zones (Fig. 5D, E). Taken together, the loss-of-function phenotypes, MD simulations and localization data support a direct function of CLCC1 in nuclear membrane fusion; a function that is sustained by Torsins.

## Discussion

How the INM and ONM are fused to complete interphase NPC biogenesis has remained a major unresolved enigma in cell biology. We show that this membrane fusion process at the NE requires the conserved membrane protein dClcc1/CLCC1, which we have identified as a Torsin interaction partner. The absence of CLCC1 and dClcc1 causes NE membrane fusion defects, analogous to those caused by *dtorsin* knockout. On the other hand, ectopic expression of dClcc1/CLCC1 is sufficient to rescue NPC biogenesis and developmental defects associated with Torsin-loss-of-function, indicating that Torsins are primarily required to sustain CLCC1 functionality. Since CLCC1 accumulates at membrane fusion sites (Fig. 5D), it emerges as the prime candidate for the elusive fusogen of the INM and ONM.

### CLCC1 as a new Torsin interaction partner

CLCC1 was identified as a novel Torsin interaction partner in this study through proximity labelling of human Torsin1A. Notably, the part of CLCC1 between TM2 and TM3 is structurally homologous to the membrane proteins Brl1 and Brr6, depletion of which leads to the formation of NE herniations and defective NPC assembly in yeast (Kralt et al., 2022, Mathiowetz et al., 2024, Mondal et al., 2025, Zhang et al., 2018), akin to the phenotypes observed upon loss-of-function of Torsins or CLCC1. In contrast, the N-terminal domain of CLCC1 involved in Torsin interaction is only present in metazoan CLCC1-like factors, an observation that supports the functional significance of the co-occurrence of the Torsin and CLCC1 protein families in this clade (Fig. S5).

The structural model of the CLCC1-Torsin complex indicates that the interaction between dTorsin and dClcc1 is established by several salt bridges and involves a three-helical bundle in the N-terminal region of dClcc1/CLCC1. We show that mutation of the three charged residues in dTorsin forming these salt bridges abolishes dTorsin function, highlighting the functional importance of Torsin-CLCC1 interaction. Notably, one of the salt bridges in the interface between human Tor1A and CLCC1 involves E302 of Tor1A, which is one of two consecutive glutamate residues that is deleted in the dystonia-causing Torsin ΔE mutation (Fig. S6F). This interface also houses a conserved glycine residue (G256 in dTorsin, G251 in human Torsin1A) that we show to be required for dTorsin functionality during *Drosophila* germ cell development.

Based on our data, we propose that defects in the interplay between Torsin and CLCC1 contribute to the cellular phenotypes and pathological outcomes associated with known loss-of-function and disease-causing Torsin mutations.

### A lipid-centric model of the membrane fusion mechanism

Modelling suggests that CLCC1 forms membrane-embedded, ring-shaped structures housing a central pore (Fig. 5, Fig. S6D; (Dai et al., 2024, Fischer et al., 2025, Guo et al., 2023, Mathiowetz et al., 2024, Wu et al., 2024)). Based on our localisation data we propose that CLCC1 assembles into oligomers at designated INM and ONM fusion sites where the membranes are bulged and closely juxtaposed. While the precise mechanism of targeting of CLCC1 to these membrane fusion sites is yet to be determined, its preference for highly curved membranes may contribute (Wu et al., 2024). Our modelling and simulations indeed demonstrate that in these CLCC1 rings, the protruding aromatic amphipathic helices (AH1) are ideally positioned to insert into the opposing membrane of the protruding INM bleb (Fig. 5A, C, F; Figs. S6D), where they may promote lipid bilayer distortion to facilitate fusion with the ONM. Interestingly, mutation of this helix elicits a dominant-negative phenotype on NPC biogenesis (Fig. 4F) and in the oligomeric state, these residues may act collectively to perturb lipid order.

Furthermore, although structurally distinct, these donut-shaped structures distantly resemble seipin rings, in the center of which bilayer remodeling takes place together with neutral lipid accumulation during lipid droplet biogenesis (Arlt et al., 2022, Klug et al., 2024, Sui et al., 2018, Walther et al., 2023, Zoni et al., 2021). In the ring-shaped CLCC1 oligomer, AH2 and TM3 are predicted to form intercalated rings of α-helices lining the interior of the central pore (Fig. S6D), where they might form a channel suited for lipid bilayer remodelling, akin to seipin rings. We thus propose that CLCC1’s suggested role in lipid flux (Mathiowetz et al., 2024, Wu et al., 2024) might be linked to its function in membrane fusion, providing a unifying view of its mode-of-action.

### Regulation of CLCC1 by Torsins for membrane fusion and lipid secretion

But what is the significance of the molecular interaction between CLCC1 and Torsins? Given that both Torsins and CLCC1 localize to sites of juxtaposed membranes over stalled NPCs assembly intermediates, we propose a direct role of CLCC1 in conjunction with Torsin in nuclear membrane fusion during NPC biogenesis. We suggest two non-exclusive models of how Torsins regulate CLCC1 function (Fig. 5F). In the first model, we posit that Torsins serve as chaperones to escort CLCC1 during its trafficking to the NE membrane fusion sites. In the second model, we put forward that Torsins dissociate CLCC1 oligomers for the completion of membrane fusion and/or recycling of subunits (Fig. 5F), the latter reminiscent of the role of NSF in post-fusion SNARE complex dissociation (Rothman, 1994, Ryu et al., 2015). In both models, CLCC1 molecules are predicted to become limiting upon Torsin loss-of-function, either because they are not efficiently delivered to the fusion site or not recycled after fusion. Thus, cells that are critically dependent on interphase NPC biogenesis, such as rapidly growing post-mitotic *Drosophila* spermatocytes and developing human neurons, are expected to be severely affected by the CLCC1 shortfall upon Torsin loss-of-function. Consistently, providing excess CLCC1 renders Torsin dispensable and suppresses the loss-of-function phenotypes in both *Drosophila* germ cells and human somatic cells.

Interestingly, loss of CLCC1 or the CLCC1-associated phospholipid scramblase TMEM41B causes ER-associated defects in the biogenesis and secretion of very-low-density lipoprotein (VLDL) particles, manifesting in ER-associated lipid droplets and hepatic steatosis (fatty liver) in mouse models (Mathiowetz et al., 2024, Wu et al., 2024). This phenocopies Torsin1A knockout in the mouse liver and the *Drosophila* fat body, a liver-like organ in flies (Grillet et al., 2016, Shin et al., 2019). We propose that CLCC1, supported by Torsins, functions in membrane remodelling for both lipid loading of VLDLs and INM-ONM membrane fusion, with the central pore of CLCC1 promoting lipid projections out of the lipid bilayer into nascent VLDL or the opposing NE membrane, respectively, perhaps supported by the activity of TMEM41B (Fig. 5F).

Finally, DYT1 primary dystonia, caused by mutations in Torsin1A, is currently an incurable disease. We show that increased expression of dClcc1/CLCC1 is sufficient to suppress the disease-like pathologies in *Drosophila* germ cells and cultured human somatic cells lacking Torsin function. These findings identify CLCC1 as a potential therapeutic target for early-onset dystonia. CLCC1 supplementation in mouse dystonia models as well as insights into the molecular mechanisms that drive CLCC1 synthesis and decay will be important steps to inform future strategies for therapeutic intervention.

## Materials and Methods

### Antibodies

For immunofluorescence and immunoblotting analyses, the following antibodies were used: rat monoclonal anti-Vasa (1:100, Developmental Studies Hybridoma Bank (DSHB, AB_760351), mouse monoclonal mAb414/anti-FG-Nup (1:100, Abcam ab24609), mouse monoclonal anti-Lamin Dm0 (1:100, DSHB, ADL84.12), anti-dELYS (gift of M. Capelson, San Diego State University), mouse monoclonal anti-V5 (1:1000, Bio-Rad, MCA1360), mouse monoclonal anti-MLF2 (1:100, Santa Cruz, sc-166874), rabbit polyclonal anti-CLCC1 (1:200, Atlas Antibodies, HPA009087), mouse monoclonal anti-ubiquitin (1:200, Enzo Life Sciences, ENZ-ABS840-0500), mouse monoclonal anti-actin (1:1000, Santa Cruz, sc-47778), anti-mouse Alexa Fluor 488 (Invitrogen, A11001), anti-rat Alexa Fluor 568 (Invitrogen A78946), anti-mouse Alexa Fluor 594 (Invitrogen, A11005), anti-mouse Alexa Fluor 647 (Invitrogen A31571), anti-rabbit Alexa Fluor Plus 800 (1:10000, Invitrogen, A32735) and anti-mouse Alexa Fluor Plus 680 (1:10000, Invitrogen, A21058).

### Molecular cloning

For the *UAS-dTorsin-GFP* wild-type constructs and mutated variants, the wildtype *dTorsin* and *dClcc1* (CG12945) ORFs were amplified from Sf9 cell cDNA, the mutant variants were generated either by gene synthesis or PCR-based site-directed mutagenesis and subcloned into a pUASt-attB-dsRed vector with a 3xP3-DsRed selection cassette (Jagannathan lab) using the XhoI and XbaI restriction sites. EGFP was PCR-amplified from pcDNA5/FRT/TO/Tor1B-EGFP (Luithle et al., 2020) and added to the C-terminus of *dTorsin* ORF using the XbaI restriction site.

The ORF encoding for human CLCC1 was amplified from human cDNA by PCR and inserted into pcDNA5/FRT/TO vector (Invitrogen) including a C-terminal GFP-tag using EcoRV and NotI cloning sites. DNA fragments containing CLCC1 mutants were ordered from TWIST Bioscience and inserted into pcDNA5 backbones with a C-terminal GFP linearized with EcoRV and NotI using Gibson assembly.

For SUN2-Venus, the human SUN2 ORF (Q9UH99-1) was PCR-amplified from SUN2-GFP (Turgay et al., 2010) and inserted into a pIRES backbone using NheI and EcoRI. Venus was amplified and inserted C-terminally using EcoRI and NotI sites. For KASH2-mTFP1, human Nesprin-2 (Q8WXH0-1, 6556-6858 aa) was PCR-amplified and inserted into a pIRES backbone using NheI and BamHI, following a 3xHA tag. mTFP1 was inserted between the transmembrane segment and luminal domain of Nesprin2 (KASH2) using BamHI and NotI sites, following insertion of the KASH2 peptide (aa 6857-6885) using NotI sites. For Venus-CLCC1, a DNA fragment encoding for human CLCC1, containing Venus-encoding sequence, behind the signal peptide (1-18 aa), was ordered from TWIST Bioscience and inserted into a pIRES backbone linearized using NheI and NotI by Gibson assembly. For Torsin1A-mTFP, Torsin1A was amplified by PCR from vectors generated in (Luithle et al., 2020) and inserted into a pIRES vector containing C-terminal mTFP1 linearized with EcoRV and BamHI by Gibson assembly. For the Torsin1A-TurboID-V5 construct, TurboID was first amplified (Addgene plasmid #107177 (Branon et al., 2018)) and cloned together with a V5-tag amplified by overlap extension PCR into the NotI/ApaI-linearized pcDNA5/FRT/TO vector (Invitrogen) by Gibson assembly. Next, the coding region of full-length human Torsin1A was cloned into the BamHI and NotI sites of pcDNA5/FRT/TO/TurboID-V5. For the SS-V5-TurboID-GFP construct (negative control), the first 69 bp of the Torsin1A coding region (signal sequence, SS), the V5 tag amplified by overlap extension PCR and the TurboID tag were inserted by Gibson assembly into the ApaI and HindIII sites of the pcDNA5/FRT/TO vector (Invitrogen). Next, using overlap extension PCR, the first 69 bp of the Torsin1A coding region encoding the signal sequence and the EGFP coding sequence including a region for a KDEL peptide was amplified by PCR and cloned into the KpnI and XhoI sites.

### Fly husbandry and strains

Flies were kept on standard Bloomington medium at 25°C, unless mentioned otherwise. *dtorsin^KO13^*, *dtorsin^KO78^* and *UASt-dtorsin* (referred as *Rescue^gene^* in this study) strains were kind gifts from the Ito lab (Wakabayashi-Ito et al., 2011). *GFP- Nup107* (BDSC35514), *Nup58-EGFP* (BDSC91768), *EGFP-Nup358* (BDSC95390), *UAS-mCherry^RNAi^* (BDSC35785*), UAS-vasa^RNAi^* (BDSC34950), *Dp(1;3)DC472* (BDSC32303), C(1)RM/C(1;Y)6 (BDSC9460), *hs-FLP* (BDSC7), *FRT82B* (BDSC86313), Ubi-NLS-RFP (BDSC30555), CG12945^EP3613^ (BDSC17145) and *UASt-NLS-GFP* (BDSC4775) were acquired from the Bloomington Drosophila Stock Center. *bam-GAL4* has been previously used (Chen & McKearin, 2003). *nos-GAL4^+VP16^* (second chromosome) was a gift from Y. Yamashita (Whitehead Institute, MIT). To generate mosaic flies containing clones homozygous for *CG12945/dClcc1^EP3613^*or *CG12945/dClcc1^KO^*, *hs-FLP*; *FRT82B, Ubi-NLS-RFP* females were crossed to either *EGFP-Nup358; FRT82B, dClcc1^EP3613^* or *FRT82B, dClcc1^KO^* males. The progeny was heat-shocked for 1 h at 37°C 2 days post egg laying. Testes from adult flies were dissected and *dClcc1^EP3613^*or *dClcc1^KO^* homozygous clones identified by lack of nuclear RFP signal. Transgenic flies were generated using phic31-mediated recombination (BestGene Inc.).

### Generation of CRISPR/Cas9 knockouts (KO) for *Drosophila* strains

*CG14103/TorIP* and *CG12945/dClcc1* KO fly strains were generated through CRISPR-mediated homology-directed repair, replacing the protein coding sequence with an attP flanked 3xP3-DsRed selection cassette, as previously described (Chavan et al., 2024). Briefly, homology arms (HA) were amplified from the 5’ and 3’UTRs (670 - 1000 bp) of *TorIP* and *dClcc1* by PCR from *yw* genomic DNA and cloned into the pBSK-attP-DsRed-attP vector using Esp3I and LguI restriction sites, respectively. Two guide RNAs (gRNAs) per gene targeting the start and stop codon were cloned into the pU6-Bbs1-ChiRNA plasmid. *TorIP* - gRNA1 5’-GTGGCGAGACGACCACTGCA-3’, gRNA2 5’-GTCTTTCAGTTTAACTCAAA-3’. *dClcc1* – gRNA1 5’- GTCTCAATGTAATCACTCCT-3’, gRNA2 5’-GGAAGATGAAGGATTCACGC-3’. The donor and gRNA plasmids were microinjected into *yw; nos-Cas9 (attP40) Drosophila melanogaster* strains and knockouts were identified based on DsRed expression (BestGene Inc). The correct integration of the DsRed cassette was verified by PCR centered around the cut site using the following primers at the *TorIP^KO^* ORF flanking regions (5’- CCGGCGTCTACCAATTGGCTG-3’ and 5’- GATCGGCCGACGTAAGCCCTA-3’) , the *dClcc1^KO^* ORF flanking regions (5’ - GTAAATCCGGAAATCTCGGCCTTC-3’ and 5’- CAGCCGCCAACACAAATCAGATG-3’) and within the DsRed cassette (5’- GATCCACAAGGCCCTGAAGC-3’ and 5’- GTACTGGAACTGGGGGGACAG-3’).

### Fertility assays

Two *yw* virgins were crossed to a single male (male fertility assay) or two *yw* males were crossed to a single virgin of the desired genotype (female fertility assay) for six days and the number of eclosed progeny was counted for 18 days post setup.

### Mammalian cell culture, transfection and polyclonal cell line generation

HCT116 and HeLa cells were cultured in Dulbecco’s Modified Eagle Medium (DMEM; Gibco) supplemented with 10% fetal calf serum (FCS; Eurobio Scientific) and 100 µg/ml penicillin/streptomycin (Corning) at 37°C with 5% CO_2_.

HeLa and HCT116 cells were transiently transfected in a 6-well plate using a mixture of 100 µl of JetPrime buffer, 3 µl JetPrime reagent (Polypus) and 1 µg of DNA per well, incubated at RT for 15 min before addition to cells. Media was changed 6 h after transfection.

To generate tetracycline-inducible cell lines, HeLa FRT/TO cells were transfected with Flp recombinase (pOG44 plasmid, Invitrogen) and pcDNA5/FRT/TO vectors encoding TurboID-tagged proteins. For each well of a 6-well plate, a transfection mixture of 200 µl of jetPRIME buffer, 3 µl jetPRIME reagent (Polypus) and 1 µg plasmid DNA (0.9 µg pOG44 and 0.1 µg pcDNA5/FRT/TO) was incubated for 15 min before being added to cells. After 48 h, cells were subjected to selection with 400 µg/ml hygromycin (Invitrogen) until individual clones were visible (2.5 - 3 weeks).

### Generation of human CRISPR/Cas9 knockout cell lines

To generate *CLCC1* knockout cell lines, CRISPR gRNAs targeting the regions around the start and stop codons of the human *CLCC1* gene were designed using the web-based tools CHOPCHOP and CRISPOR (Concordet & Haeussler, 2018, Montague et al., 2014). gRNA1 5’-AGTGTTATCTTAACTCTAGA-3’ and gRNA2 5’-GTATACTTTACAAAGCTCGA-3’ were selected to target the regions upstream of the start codon (∼ 400 bp) and downstream of the stop codon (∼ 40 bp), respectively. Forward and reverse DNA oligonucleotides (Merck) containing the regions encoding for the gRNAs with appropriate overhangs were annealed and ligated into the BsaI-digested vector pC2P-Cas9 (Welte et al., 2019). HeLa or HCT116 cells were transfected with 1 µg of each plasmid DNA using JetPRIME (Polypus). After 24 h, cells were transferred to 10 cm plates and selected with 2 µg/ml puromycin for 48 h. After 2.5 - 3 weeks, individual colonies were isolated, expanded into cell lines and characterized by PCR. To confirm full deletion of the CLCC1 coding sequence, the following primers were used: forward 5’-CCTAGCAATGGAATTGCTGGG and reverse 5’- CAAAGAGAAGATGAAATGAAAAGC.

To generate *TOR1AIP1/TOR1AIP2* double knockout HeLa cell lines, a similar procedure was applied. First, a complete knockout of the *TOR1AIP2* gene was generated using gRNA1 5’-AGCATGAGTCTGTTGGGACT-3’ and gRNA2 5’- CTACCTAAATACTACATGTA-3’ targeting gene regions upstream of the start codon (∼ 90 bp) and downstream of the stop codon (∼ 590 bp), respectively. Clones were characterized by PCR (forward primer: 5’- TGCCAGTAGACACAGTTTGAAAATC-3’; reverse primer: 5’- GTTTGTGGGAACCTCCTGTGCATTG-3’). A confirmed*TOR1AIP2* KO clone was subjected to knockout generation of the *TOR1AIP1* gene using gRNA1 5’-TCAACAACTATGGCGGGCGA-3’ and gRNA2 5’-AGTGAGAAATACGGCTCCAG-3’ targeting gene regions downstream of the start codon (∼ 5 bp) and upstream of the stop codon (∼ 60 bp), respectively. *TOR1AIP2/TOR1AIP1* knockout clones were characterized by immunoblotting and by PCR. The double deletion of the *TOR1AIP1/TOR1AIP2* genes (which lie adjacent to each other, divergent orientation) was confirmed by PCR (forward primer: 5’-TGCCAGTAGACACAGTTTGAAAATC-3;’ reverse primer 5’- AGGCAGAAGCTAAGGAACTCACACC-3’). Although the DKO clone seemed initially complete by PCR analysis, MS later revealed the presence of some LAP1 peptides. Phenotypically, the cells showed the expected phenotypes, similar to what has been reported (Laudermilch et al., 2016).

### Immunofluorescence staining and microscopy

Seven to nine testes or four to five ovaries per condition were dissected in 1x PBS, fixed in 4% formaldehyde in 1x PBS for 20 min and washed with 0.1% Triton X-100 in 1x PBS (1x PBS-T) for at least 2 h at room temperature (RT) on a rotator. Samples were incubated in primary antibodies diluted in 3% bovine serum albumin (BSA) in 1x PBS-T overnight at 4°C. Next day, after 1 h of washing in 1x PBS-T, tissues were incubated with secondary antibodies diluted in 3% BSA in 1x PBS-T overnight at 4°C. After 1 h of washing in 1x PBS-T, testes and ovaries were mounted in VECTASHIELD medium with DAPI (Vector Laboratories) and imaged with a Zeiss LSM 780 upright Confocal Microscope with 63x 1.4NA Oil Plan-Apochromat objective operated by the ZEN software. Images were analysed by ImageJ2 and Imaris (for 3D reconstruction).

For immunofluorescence of human cells, cells on coverslips were fixed in 4% paraformaldehyde (PFA) in 1x PBS for 10 min at RT, washed thrice with 1x PBS, permeabilized with 0.2% Triton X-100 and 0.02% SDS in 1x PBS for 5 min at RT, washed as above and blocked in 2% BSA in 1x PBS (blocking solution) for at least 20 min at RT. Next, cells were incubated with primary antibodies diluted in 2% BSA in 1x PBS for 2 h at RT, then washed twice with blocking solution and incubated with secondary antibodies diluted 1:300 in blocking solution for 1 h at RT. Then, coverslips were washed thrice with 1x PBS, incubated in 1 µg/ml Hoechst for 10 min at RT and washed twice. Lastly, coverslips were mounted in VECTASHIELD medium (Vector Laboratories) and imaged with a Zeiss LSM 780 upright Confocal Microscope with 63x 1.4NA Oil Plan-Apochromat objective operated by ZEN software. Images were analysed using ImageJ2 (version 2.16.0/1.54p; http://imagej.net; National Institutes of Health, USA), and graphs were generated using GraphPad Prism (version 10.5.0; GraphPad Software, San Diego, CA, USA).

To quantity ubiquitin or MLF2 foci, cells were binned into predefined categories: strong foci, weak foci, or no foci and counted. For rescue experiments with GFP-tagged constructs, only GFP-poistive cells were considered.

### RNA fluorescence *in situ* hybridization (FISH)

RNA in situ hybridization in *Drosophila* tissues was performed as previously described (Chavan et al., 2024). Briefly, seven to nine testes or four to five ovaries per condition were dissected in RNase-free PBS and fixed in 4% formaldehyde in RNase-free PBS for 30 min. After three washes with PBS for 5 min, tissues were dehydrated in RNase-free 70% ethanol overnight at 4°C on a nutator. Next day, tissues were treated with wash buffer (2x SSC, 10% deionized formamide) for 5 min at RT before incubation in freshly made hybridization mix (2x SSC, 10% dextran sulfate, 1 g/L *E. coli* tRNA, 2 mM vanadyl ribonucleoside complex, 0.5% RNase-free BSA, 10% deionized formamide, 125 nM oligo(dT)-Cy5 (Microsynth) probe) overnight at RT on a nutator. The next day, samples were washed twice with wash buffer for 30 min at RT on a nutator and mounted with VECTASHIELD medium with DAPI (Vector Laboratories). Imaging was done with a Zeiss LSM 780 upright Confocal Microscope with 63x 1.4NA Oil Plan-Apochromat objective operated by the ZEN software.

### Förster Resonance Energy Transfer (FRET)

HeLa *CLCC1* KO cells were transiently co-transfected with plasmids encoding mTFP1- and Venus-tagged proteins in Ibidi imaging chambers, using 0.5 µg DNA per plasmid and JetPrime. Cells were fixed with 4% PFA for 10 min 24 h after transfection, and washed 6 times with 1x PBS. A Zeiss LSM 880 inverted confocal microscope (equipped with a 63x 1.4 NA oil objective and operated by the Zen software) was used to carry out acceptor photobleaching. For each cell, 4 regions of interest at the nuclear envelope were designated, two underwent bleaching of the acceptor Venus (514 nm laser, 100% power, 120 iterations), two were unbleached to serve as controls for photobleaching during imaging. An additional background region was also designated and measured, away from the cell. Data was analysed in Microsoft Excel, intensity measurements of mTFP1 were background subtracted and corrected for bleaching (relative change in intensity in the unbleached NE regions calculated and applied to the post-bleaching intensity measurements). FRET efficiencies were then calculated using the following formula: FRET efficiency [%] = 100 x (I_post_ - I_pre_)/ I_post_, where I_pre_ is donor intensity before bleaching and I_post_ is donor intensity after bleaching.

### Immunoblotting

Proteins were transferred onto a 0.45 µm nitrocellulose membrane (Amersham Protran) using semi-dry blotting for 1 h at 180 mA. Membranes were blocked in 5% (w/v) non-fat dry milk in 0.1% Tween-20 in 1x PBS for 10 min, then incubated overnight at 4°C with primary antibodies diluted in blocking buffer. The following day, membranes were washed with blocking buffer three times, incubated with an appropriate secondary antibody for at least 1 h at RT, washed with 0.1% Tween-20 in 1x PBS three times and visualized using the Odyssey Imaging System (LI-COR).

### Proximity labelling *in vivo*, streptavidin pulldown and on-bead digestion

Approximately 40x 10^6^ HeLa FlpIn cells per condition were induced with 20 ng/ml tetracycline (Sigma) for 24 h and treated with 200 µM biotin (Sigma) for 10 min at 37°C and 5% CO_2_ prior to harvesting. For cell lysis, a denaturing buffer (50 mM Tris-HCL pH 7.5, 7.5 M urea) supplemented with protease inhibitors (10 µg/ml Leupeptin, 10 µg/ml Aprotinin, 1 µg/ml Pepstatin A) and 1 mM PMSF was used. The lysate was subjected to three rounds of 30 - 60 s sonication (Bandelin Sonoplus, UW50, TS102) and cleared by centrifugation at 15’000x g for 15 min at 4°C. The supernatant was incubated for at least 2 h with streptavidin sepharose beads (GE Healthcare) at 4°C. The beads were washed three times with wash buffer (50 mM Tris-HCL pH 7.5, 7.5 M urea), followed by one wash with the wash buffer including 1% Triton X-100. Co-precipitated proteins were proteolyzed on beads as previously described (Uliana et al., 2023, Wisniewski et al., 2009). Beads were transferred to Vivacon 500 centrifugal units (Sartorius, MWCO 10 kDa), treated with 8 M urea, reduced with 5 mM TCEP (tris(2-carboxyethyl)phosphine) for 30 min at 37°C, alkylated with 10 mM iodoacetamide for 30 min at 37°C and washed with 100 mM ammonium bicarbonate prior to bead digestion with 1 µg Trypsin (Promega) and 0.5 µg LysC for 14 h at 37°C. Peptides were eluted by centrifugation at 8000x g for 20 min and after quenching proteolysis with 5% formic acid, the peptides were purified using C18 cleanup columns (The Nest Group) following the manufacturer’s protocol. Subsequently, eluted peptides were dried using a SpeedVac.

### Mass spectrometry and data analysis

Prior to LC-MS/MS analysis, peptides were resuspended in 2% acetonitrile and 0.1% formic acid and then injected into an Orbitrap Q Exactive Plus mass spectrometer (Thermo Fisher) coupled to an EASY-nLC 1000 liquid chromatography system (Thermo Fisher). Peptides were separated using a 75 μm ID x 400 mm column (New Objective), in-house packed with ReproSil Gold 120 C18 (1.9 μm; Dr. Maisch GmbH) with a gradient from 3% to 25% in 170 min and 25% to 40% using buffer A (0.1% formic acid) and buffer B (80% acetonitrile, 0.1% formic acid). Samples were acquired in data-dependent acquisition (DDA) with MS1 scan (scan range = 350 - 1500, R = 70,000, maximum injection time 100 ms, and AGC target = 1e6), followed by 20 dependent MS2 scans (scan range = 120 - 2000, R = 17,500, maximum injection time 55 ms, and AGC target = 1e5). Peptides with charges between 2 and 7 were isolated (m/z = 1.8) and fragmented (NCE 25%). Acquired spectra were searched using MaxQuant (version 2.4.13) against the human proteome database (downloaded from UniProt in April 2021; 20’388 proteins) extended with reverse decoy sequences and contaminant list from MaxQuant. The search parameters were set to allow specific tryptic peptides, two missed cleavages, fixed cysteine carbamidomethylation, variable methionine oxidation and protein N-terminal acetylation. The MS and MS/MS search parameters were set to +/-10 ppm tolerance. A false discovery rate of < 1% was applied at both the PSM and protein levels. Protein abundance was determined from the intensity of the top two peptides, with the “match between runs” option enabled. Intensity values for proteins identified in all replicates in at least one condition were log_2_ transformed, median-normalized and imputed using random sampling from a normal distribution derived from the lowest 1% of values.

Protein intensities were subjected to ANOVA testing (Torsin1A vs. GFP) and p-values were corrected using the Benjamini-Hochberg procedure. Protein hits with log_2_-fold change > 2 and a p-value < 0.01 were considered for further analyses. The mass spectrometry raw data, data analysis scripts and tables generated have been deposited to the ProteomeXchange Consortium (PRIDE - PRoteomics IDEntifications Database).

### Transmission Electron Microscopy (TEM)

Seven to nine testes per condition were dissected in freshly prepared fixative (2.5% glutaraldehyde, 0.1 M sodium cacodylate) and stored at 4°C. The following day, samples were washed three times with 0.1 M sodium cacodylate buffer, incubated in 1% osmium tetroxide (0.1 M sodium cacodylate) for 1 h, washed again, and incubated overnight at 4°C. The next day, samples were washed three times with ddH_2_O incubated in 1% uranyl acetate for 1 h and washed again. Dehydration was performed in a series of ethanol solutions (70% (v/v), 80%, 96% and 100%) with two 10 min washes per step. Samples were then incubated in 25% Epon/Araldite in propylene oxide for 30 min, followed by 50% Epon/Araldite overnight incubation at RT. The next day, samples were incubated in 100% Epon/Araldite for 30 min at RT and polymerized at 60°C for 30 h. Ultrathin sections (70 nm) were prepared using an ultramicrotome, post-stained with lead citrate and examined with a Talos 120 transmission electron microscope at an acceleration voltage of 120 KV using a bottom mounted Ceta camera and the MAPS software for automatic image acquisition (Thermo Fisher Scientific, Eindhoven, The Netherlands). Images were acquired at 1.7 nm per pixel. TEM was conducted at the Center for Microscopy and Image Analysis (University of Zurich). Image analysis was performed using MAPS and ImageJ.

### Immunogold Electron Microscopy (TEM)

Cells grown on 15 mm round glass coverslips were fixed for 10 min in a fixative solution containing 4% PFA (electron microscopy-grade) and 0.05% glutaraldehyde (0.2 M HEPES buffer pH 7.4) at RT, then further fixed in 4% PFA (0.2 M HEPES buffer) for 50 min at RT. After three washes with 1x PBS, cells were incubated in 50 mM glycine for 10 min, permeabilized in 0.25% saponin in 0.1% BSA, 1x PBS for 10 min at RT, and then blocked in blocking buffer (5% goat serum, 0.2% BSA, 0.1% saponin, 50 mM NH_4_Cl, 20 mM phosphate buffer, 150 mM NaCl) for 1 h at RT. Following blocking, cells were incubated with the anti-GFP primary antibody (A-11122, Thermo Fisher) for 90 min at RT, washed with 0.1% saponin in 0.1% BSA, 1x PBS and incubated with nanogold-conjugated secondary antibodies (Nanoprobes) at RT for 1 h. The samples were then fixed with 1% glutaraldehyde for 30 min, followed by treatment with gold enhancement solution (Nanoprobes) according to the manufacturer’s instructions. Next, the samples were post-fixed with reduced osmium (1% OsO_4_, 1.5% potassium ferrocyanide in 0.1 M cacodylate buffer, pH 7.4) for 1 h on ice. After several washes in Milli-Q water, sections were incubated in 0.5% uranyl acetate at 4 °C overnight. Samples were dehydrated in a graded ethanol series, embedded in epoxy resin and polymerized at 60 °C for 48 h. Ultrathin serial 70 nm thick sections were obtained using an ultramicrotome (UC7, Leica Microsystems), collected on formvar/carbon supported copper grids, stained with uranyl acetate and Sato’s lead solution. Images were acquired using a Talos L120C Transmission Electron Microscope equipped with a Ceta CCD camera (FEI, Thermo Fisher Scientific) operating at 120 kV.

### Evolutionary Analyses

#### CLCC1 distribution across Opisthokonta

To map the phylogenetic distribution of CLCC1 within Holozoa (the clade containing animals and their ancestors), a dataset of 121 proteomes was constructed using UniProt data (UniProt, 2025). These proteomes were selected to adequately cover the known taxonomic diversity of Metazoa. Since the quality of the proteome can affect the subsequent analysis using custom-built Hidden Markov Models (HMMs), their quality was assessed using BUSCO v5.7.1 (Manni et al., 2021) with the eukaryota lineage of 255 markers. Starting from the human CLCC1 sequence (UniProt ID Q96S66), we used the best 1-domain score from phmmer from the HMMER package (version 3.4) (Eddy, 2011) to retrieve the top-scoring hit from each proteome. From this set, we rejected those hits that, when used as a query for a phmmer search against the human proteome, did not return Q96S88 as the top-scoring hit. The orthology of these best ‘bi-directional’ hits to be descended from a common ancestral protein was verified by constructing maximum-likelihood gene trees using MAFFTv7.505 with the E-INS-i option enabled (Katoh & Standley, 2013) for alignment. The phylogenetically noisy and uninformative positions were removed using TrimAl v1.4.rev15 build[2013-12-17] (Capella-Gutierrez et al., 2009) with the -gappyout option enabled. Since this option removed over 70% of the positions in the alignment, the trimming was redone with more relaxed parameters, only removing those positions/columns in the alignment composed of over 40% gaps. To avoid potential artefacts in the gene tree, we further removed those sequences that were composed of more than 70% gaps. Gene trees were constructed using FastTree Version 2.1.11 Double precision (No SSE3) (Price et al., 2010) and visualized with FigTree 1.4.4 [https://tree.bio.ed.ac.uk/software/figtree/]. Outliers that either had very long branches or were badly aligned in the trimmed multiple sequence alignment were manually removed.

This trimmed alignment was used to build an HMM with hmmbuild under default options from HMMER to then iteratively search each of the 121 selected proteomes. In each iteration, we selected all hits in each species that had a best 1-domain E-value lower than 1E-5. If there was no hit in a certain species, the top-scoring hit in terms of the best 1-domain score was selected. The orthology among selected sequences was verified as described above. All verified orthologs in each iteration were used to build the HMM for the next iteration. We stopped after two iterations since the HMM searches began to retrieve non-specific hits, yielding 429 hits. To remove outliers, we used InterProScan 5.73-104.0 (Jones et al., 2014) with the SMART (Letunic et al., 2012), PANTHER (Thomas et al., 2022), SUPERFAMILY (de Lima Morais et al., 2011), Pfam (Paysan-Lafosse et al., 2025) analyses enabled, and removed every sequence with a PANTHER annotation different from the human CLCC1 protein (PTHR34093 or ‘CHLORIDE CHANNEL CLIC-LIKE PROTEIN 1’). However, if a sequence had no annotation, we retained the sequence. This step removed 127 sequences, leaving 302 sequences. Maximum-likelihood gene trees were constructed using IQ-TREE multicore version 2.3.6 (Minh et al., 2020) with 1000 ultrafast bootstraps (Hoang et al., 2018). The best-fitting model was selected by using the Bayesian information criterion scores from IQ-TREE’s ModelFinder, including three mixture models (LG+C20,LG+C40,LG+C60) for testing. We removed outliers and visualized the profile using iToL (Letunic & Bork, 2024).

To extend this phylogenetic profile to all Opisthokonta (the clade containing animals, fungi, and their deeper-branching ancestors), we assembled a set of 74 Opisthokont proteomes taken from UniProt, EukProt (Richter et al., 2022) and other sources (Tables S2, S3) and assessed their quality using BUSCO v5.7.1 (Manni et al., 2021).

The trimmed alignment from the second HMM search iteration for CLCC1 was used to search these proteomes, and all hits were selected with a best 1-domain E-value lower than 1E-5. All hits without the PANTHER annotation PTHR34093 removed, while hits with no annotation were retained. The orthology of the remaining hits was verified as described above and the profile visualized using iToL (Letunic & Bork, 2024). An interactive version of the tree can be found at https://itol.embl.de/tree/101162135313091760269782.

### Distribution of Torsins across Opisthokonta

The phylogenetic profile of the Torsin family across Opisthokonta was constructed as described above. Briefly, we selected the human TOR1A (UniProt ID O14656) and retrieved the best, bidirectional hits for this protein across the set of 74 Opisthokont proteomes. After verifying the orthology of these hits, the trimmed alignment was used to construct an HMM and search each of the selected proteomes. We selected all hits that had a best 1-domain E-value lower than 1E-7 in each species. In species with no hits with an E-value below this threshold, we selected the best scoring hit. From this set which contained both Torsins and closely related proteins, we selected the hits belonging specifically to the Torsin family by identifying the monophyletic clade containing the human and fruit fly Torsins. We rebuilt gene trees using IQ-TREE (Minh et al., 2020) for this subset and narrowed our selection again to monophyletic clades containing the human TOR1A and fruit fly dTorsin. We used this to construct a phylogenetic profile for the TOR1A family alongside other Torsins.

### MD simulations

For simulations of CLCC1, a 16-mer structure of CLCC1 (uniport ID: Q96S66) was predicted using the AlphaFold3 web server (Abramson et al., 2024). All coarse-grained (CG) simulations were performed with GROMACS (Abraham et al., 2015) v.2023.3, using the Martini 3 force field (Souza et al., 2021) using a similar protocol as described in our previous work (Fischer et al., 2025). To convert the protein structure predictions from all-atom to CG resolution, the Martinize2 script (Kroon et al., 2022) was used. An elastic network with a force constant of 500 kJ/mol·nm² was applied to preserve the secondary structure of the protein, merging all individual subunits of the protein complex. Next, the insane.py script (Wassenaar et al., 2015) was used to generate a lipid bilayer with the following composition: DOPC: DOPE: DOPS: DOPA: DODG (69:20:5:2:4). The bilayer was equilibrated using a five-step protocol that progressively increased the time step while gradually reducing the position restraints on the lipid headgroups, ultimately removing all restraints in the final step to allow the lipids to move freely. The temperature (310 K) and pressure (1 bar) were maintained using the Berendsen thermostat and barostat (Berendsen et al., 1984), respectively. The system was then stripped of solvent and ions to obtain only the equilibrated bilayer which was used in subsequent simulations. For vesicle simulations, we utilized the Martini Maker tool of CHARMM-GUI (Qi et al., 2015) to generate a 12 nm-radius vesicle made solely of DOPC. The vesicle was equilibrated following the CHARMM-GUI recommended five-step equilibration approach, with simulation durations of 100, 50, 25, and 25 ns, respectively. The time-steps and position restraints were treated as previously described.

To create a model system of the protein embedded in a planar bilayer, the protein was inserted into the equilibrated bilayer, removing all lipids within 0.7 nm of the protein to prevent steric clashes. Both the protein insertion and the lipid removal steps were executed by an in-house TCL script on VMD (Humphrey et al., 1996). As a result, several lipids remained within the inner ring of the 16-mer oligomers. The system was then solvated with water and ions to achieve charge neutrality and a final excess concentration of 0.15 M NaCl. Energy minimization was performed using the steepest descent algorithm for 5000 steps, with a position restraint of 1000 kJ/mol·nm² applied to the protein backbone beads. The system was then equilibrated using the five-step protocol, incrementally increasing the time step from 1 fs to 15 fs while steadily reducing position restraints on the protein backbone beads. The temperature (310 K) and pressure (1 bar) were maintained using the Berendsen thermostat and barostat, respectively. Water and ions were removed to obtain the equilibrated structure of the protein in the membrane, which was then used to create the prospective systems including two membranes.

For the system where the protein is in a flat bilayer opposing a vesicle, the respective equilibrated structures were positioned apart along the bilayer normal with a minimum distance of around ∼4 nm. This combined system was then sufficiently solvated, ionized, and re-equilibrated using a similar five-step protocol. Finally, four independent replicas of this system were simulated for 1.5 μs each.

For the system containing two opposing planar bilayers with proteins in each, the equilibrated membrane bilayer and protein system was duplicated, and the bilayers were positioned apart by around ∼6 nm, ensuring that proteins face each other. The entire system was then re-solvated with 0.15 M NaCl solution. To maintain the water density in the inter-bilayer space, we implemented an equilibration protocol in which a 2 nm diameter pore was temporarily created in one of the bilayers to allow the passage of water and ions from the intermembrane space to outside of membrane and vice versa. During this process, lipid tails were restrained to stay away from the pore using a flat-bottom restraint potential (Biriukov & Javanainen, 2023). The pore was then gradually filled through further equilibration steps by reducing both the diameter and the force constant for the flat bottom restraint, resulting in a stable configuration for production runs. Production simulations across four replicas continued for 1.5 μs, which was consistently found to be sufficient for lipid mixing to occur between the two bilayers.

All production simulations were performed with a time step of 20 fs. The temperature was maintained at 310 K using a V-rescale thermostat (Bussi et al., 2007) with a coupling constant of 1 ps, while the pressure was controlled at 1 bar using a Parrinello-Rahman barostat (Parrinello & Rahman, 1981) in a semi-isotropic manner, coupling the system laterally while allowing the direction parallel to the bilayer normal to fluctuate freely.

To quantify the distances between the inner and outer nuclear membranes (INM and ONM) over time, the “gromacs mindist” tool (Abraham et al., 2015) was used. Only the phosphate (PO₄) beads of lipids in both membranes were considered for these calculations. The interaction frequency of CLCC1 with the INM (Fig. 5A-C) was determined using the “gromacs select” tool (Abraham et al., 2015), specifying protein residues within 0.7 nm of INM lipids. Based on the calculated interaction frequencies, protein residues were colored in VMD (Humphrey et al., 1996), with red indicating the lowest frequency and blue indicating the highest. All graphical plots were generated using Matplotlib (Hunter, 2007), and molecular visualizations were prepared using VMD.

## Acknowledgements

We thank K. Mahan and Dr. I. Zemp for critical comments on the manuscript. We are very grateful to Dr. C. Schlieker (Yale University) for sharing the Tor4KO cell lines. We thank Drs. N. Ito (Massachusetts General Hospital, Boston) and M. Capelson (San Diego State University) for providing *dtorsin^ko^ Drosophila* lines and dELYS antibodies, respectively, Dr. A. Rothballer for the development of the SUN-KASH FRET sensor, D. Baumann for help with cell line generation, Drs. A. Chavan, F. Brändle, and H. Stocker for advice on fly experiments, B. Todorovic, U. Lüthi, C. Kaiser, N. Schilling, J. Riemann, A. Kaech and J.M. Mateos (University of Zurich) and E. Vezzoli from the ALEMBIC facility (San Raffaele Scientific Institute, Milan) for support of electron microscopy, and the staff of IBC (ETH Zurich) for technical and infrastructural support. Fluorescence microscopy was performed on instruments maintained by ScopeM (ETH Zurich), with expert support by Dr. T. Schwarz. We wish to also thank our colleagues K. Weis and his team, in particular J. Fischer, for the fruitful collaboration on the Brl1 and CLCC1 protein families.

## Data availability

Mass spectrometry data generated in this study have been deposited to PRIDE with the identifier PXD070365 and are publicly available as of the publication date.

