## Supplemental Material for "Dystonia-associated Torsins sustain CLCC1 function to promote membrane fusion of the nuclear envelope for NPC biogenesis"

**INVENTORY OF SUPPLEMENTAL MATERIAL**

- **8 Supplemental Figures**

- **4 Supplemental Tables**

  Table S1: Protein enriched on Torsin1A by TurboID (proximity labelling)

  Table S2:121 Metazoa proteomes taken from UniProt for evolutionary analysis

  Table S3: 74 Opisthokont proteomes

  Table S4: Mass spectrometric analysis of TurboID experiment (relates to Fig. 2E), Excel file

- **2 Supplemental Movies**

  Movie S1: 3D reconstruction of EGFP-Nup107 fluorescence of spermatocytes from control testes.

  Movie S2: 3D reconstruction of EGFP-Nup107 fluorescence of spermatocytes from *dtorsin*$^{KO13}$ testes.

### Supplementary figure section

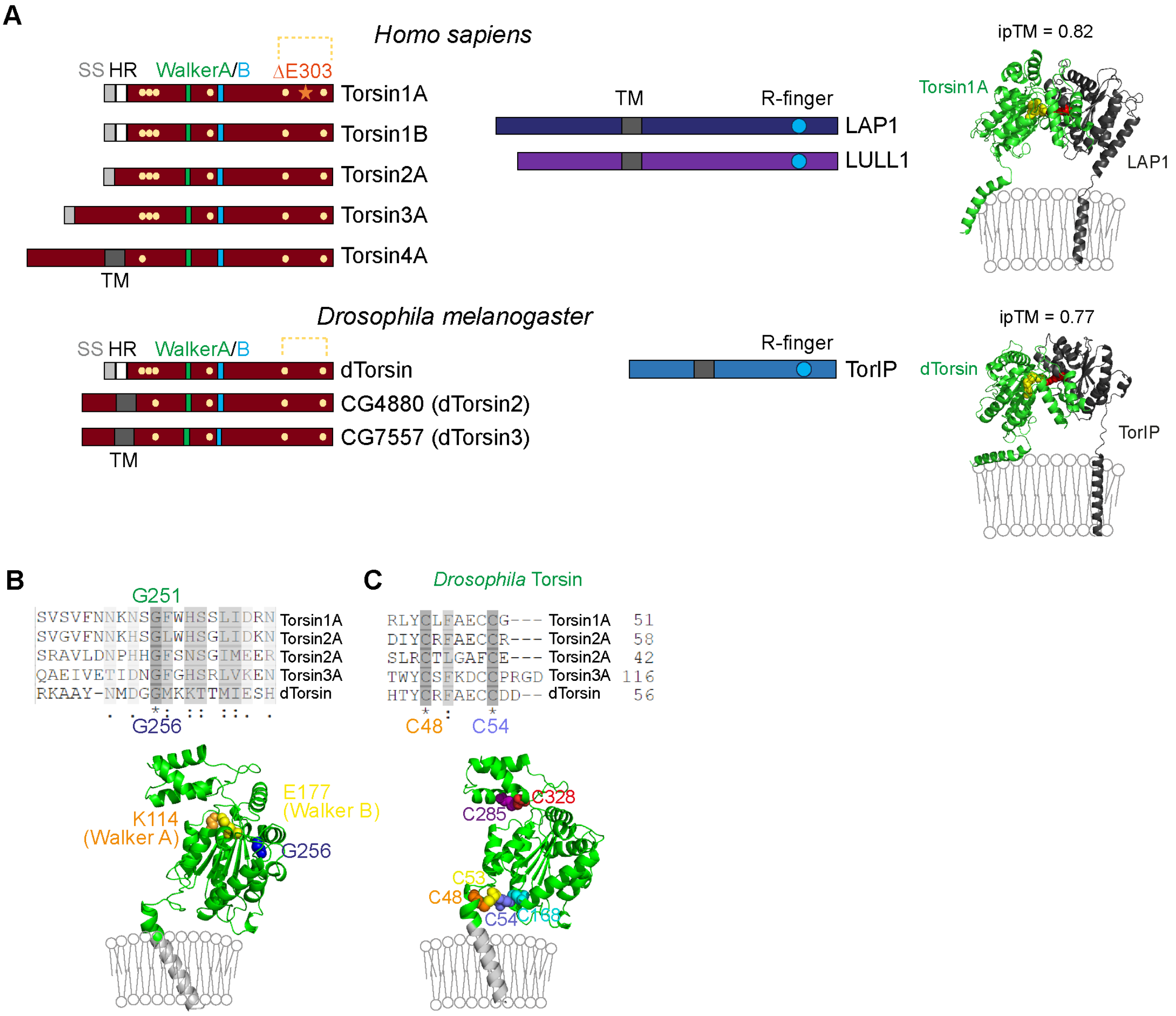


**Figure S1: The human and fly Torsin family members and their activators.**

**(A)** Schematic depiction of the five human and three fly Torsin family members, which are P-loop ATPases belonging to the AAA+ ATPase family. The Walker A (P-loop) and Walker B motifs are indicated in green and blue, respectively. Human Torsin1A, Torsin1B, Torsin2A and Torsin3A as well as *Drosophila* Torsin (dTorsin) are targeted to the ER lumen by an N-terminal signal sequence (SS). After the SS, Torsin1A, Torsin1B and fly dTorsin contain a hydrophobic region (HR) that is assumed to attach the enzymes to the inner leaflet of the ER membrane {Vander Heyden, 2011 #227}{Zhao, 2016 #10}. In *Drosophila melanogaster*, the homologous protein dTorsin (CG3024) is most closely related to human Torsin1A. The two uncharacterized transmembrane proteins CG4880 and CG7557 (dTorsin2 and dTorsin3) resembleTorsin4A. Human Torsin4A as well as Drosophila dTorsin2 and dTorsin3 are type 2 membrane proteins anchored in the ER membrane by transmembrane segments (TM). The orange asterisk in human Torsin1A marks the disease-causing deletion of the glutamate residue 303 (ΔE303). Yellow dots represent conserved cysteine residues. The dashed line shows a disulfide bond. The human and fly Torsin activators are type 2 transmembrane proteins and contain a conserved arginine residue (R-finger) important for ATPase activation of Torsins. Right: Structural models of the Torsin1A/LAP1 and dTorsin/TorIP complexes were generated using AlphaFold. The conserved arginine finger of LAP1 (R563) and TorIP (R292) (in red,) and the Walker A and Walker B motifs (K108 and E171 of Torsin1A, and K114 and E177 of dTorsin) (in yellow) are depicted in space-filling representation.

**(B)** AlphaFold3 model of *Drosophila* Torsin (F6JQA6, residues 27–340, signal peptide excluded), showing residues of Walker A and Walker B motifs as well as the conserved G256 residue in space-filling representation in the structure. The hydrophobic region (HR) is shown in light gray, associated with the membrane. Alignments were done on https://www.uniprot.org/align.

**(C)** AlphaFold3 model of *Drosophila* Torsin (F6JQA6, residues 27–340, signal peptide excluded), showing predicted disulfide bonds: N-terminal C48–C53 and C54–C168, and C-terminal C285–C328. The hydrophobic region (HR) is shown in light gray, associated with the membrane. Alignments were done on https://www.uniprot.org/align.

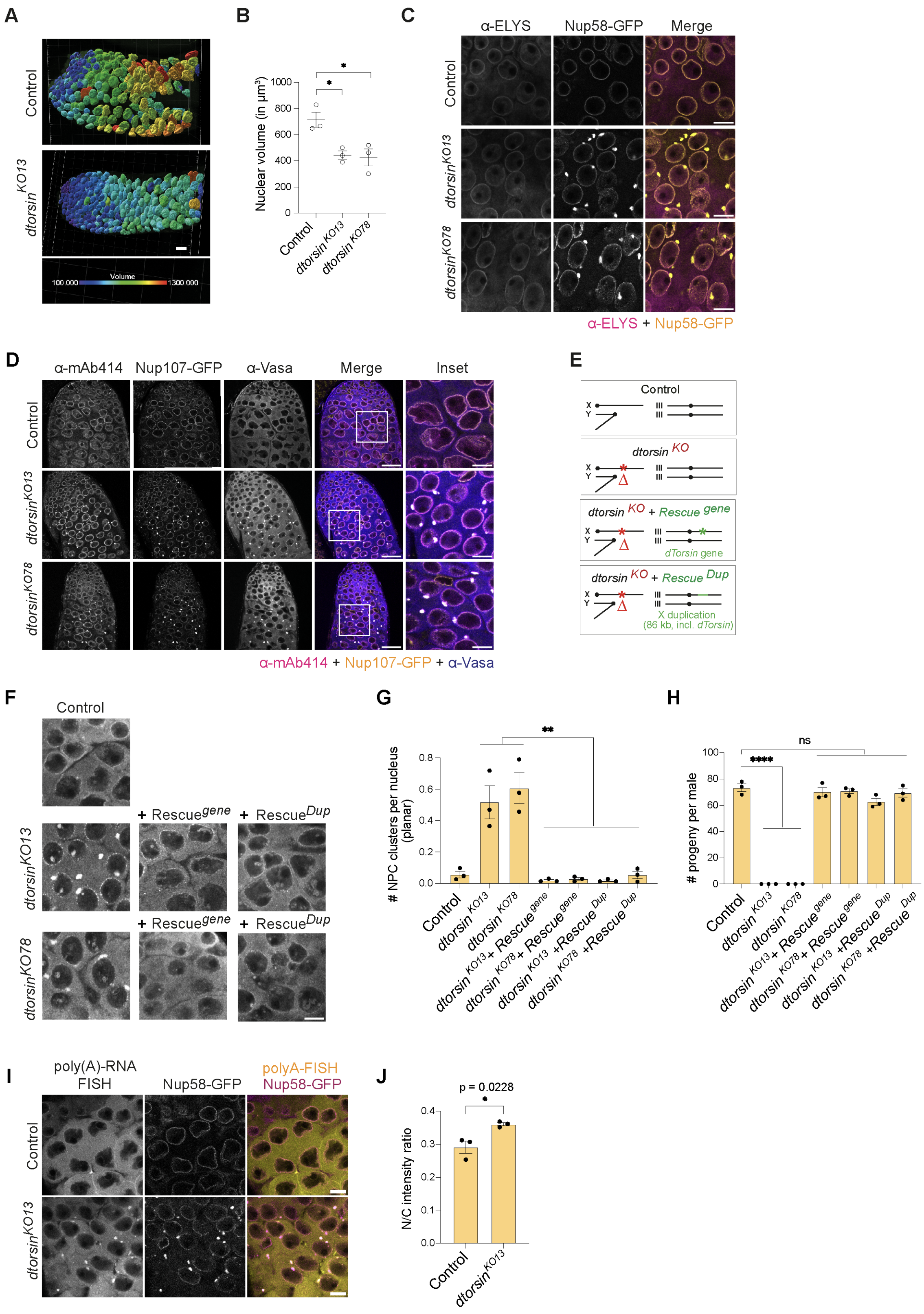

**Figure S2: *dtorsin*$^{KO}$ flies are sterile due to germline developmental arrest.**

**(A)** Visualization of nuclear volume of control and *dtorsin*$^{KO13}$ spermatocytes based on nuclear GFP signal using 3D reconstructions of confocal images (Imaris). Volume units are µm$^3$. Scale bar: 20µm.

**(B)** Nuclear volume quantification from (A) (N = 3, n = 150, * p ≤ 0.05, student's unpaired t-test, two-tailed, mean +/- SEM).

**(C)** Representative confocal images of spermatocytes from control or the two *dtorsin*$^{KO}$ strains expressing Nup58-GFP (yellow) and immunostained with anti-ELYS antibodies (magenta). Scale bars: 10 µm.

**(D)** Representative confocal images of spermatocytes from either control or the two *dtorsin*$^{KO}$ strains expressing GFP-tagged Nup107 and immunostained using mAb414 (FG-Nups, yellow) and anti-Vasa (magenta) antibodies. Scale bars: 30 µm, inset scale bars: 10 µm.

**(E)** Scheme of fly genotypes and crosses used for rescue experiments. Knockout of the endogenous *dTorsin* gene is represented by a red asterisk in the *dtorsin*$^{KO}$ strain. Rescue experiments were performed with either *bam-Gal4*-driven *UASt-Torsin* expression (Rescue$^{gene}$, green asterisk represents the location of the *UASt-Torsin* transgene) or using a *dTorsin*-containing 86 kb X;3 chromosomal duplication (Rescue$^{Dup}$, green bar represents the duplicated X segment).

**(F)** Representative confocal images of *dtorsin*$^{KO}$ spermatocytes in the absence and presence of the dTorsin wild-type rescue constructs described in (E), immunostained with anti-Vasa antibody. Scale bar: 10 µm.

**(G)** Quantification of the number of NPC/Vasa-positive foci in spermatocytes from the indicated genotypes. ($N = 3$, $n \geq 200$, *** $p \leq 0.001$, student's unpaired t-test, two-tailed, mean +/- SEM).

**(H)** Fertility assays of individual males from the indicated genotypes as described in (E). Each dot represents the number of progeny sired by a single male crossed to two wild-type virgin females ($N = 3$ independent crosses, **** $p \leq 0.0001$, ns $p > 0.05$, student's unpaired t-test, two-tailed, mean +/- SEM).

**(I)** Representative confocal images of poly(A)-RNA FISH on control and *dtorsin*$^{KO13}$ spermatocytes expressing GFP-tagged Nup58 (magenta) using oligo(dT)-cy5 probes (yellow). Scale bars: 10 µm.

**(J)** Quantification of nuclear to cytoplasmic (N/C) ratio of poly(A)-RNA signal intensity from control and *dtorsin*$^{KO13}$ spermatocytes ($N = 3$, $n \geq 150$ cells, * $p \leq 0.05$, student's unpaired t-test, two-tailed, mean +/- SEM).

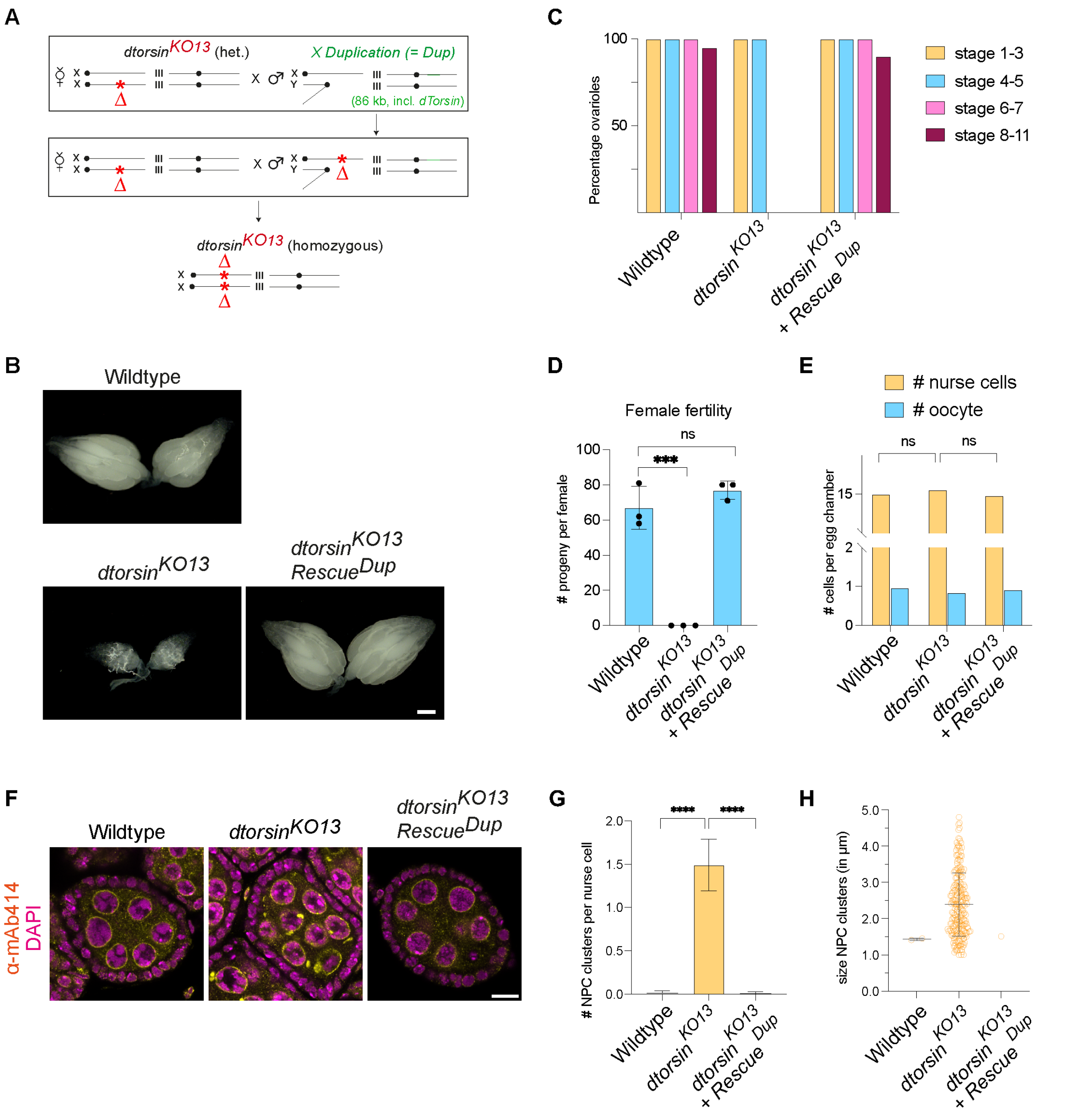


### Figure S3: Female *dtorsin*$^{KO}$ flies are sterile, akin to *dtorsin*$^{KO}$ males.

**(A)** Crossing scheme used to generate homozygous *dtorsin*$^{KO}$ female flies. *dtorsin*$^{KO}$ females are usually not observed since males carrying the X-linked *dtorsin*$^{KO}$ alleles are sterile. This obstacle was overcome by using *dtorsin*$^{KO}$ males that retain fertility due to the presence of a heterozygous X;3 *dTorsin*-containing chromosome duplication. Crossing these males to *dtorsin*$^{KO/+}$ females yields homozygous *dtorsin*$^{KO}$ females at sub-Mendelian ratios.

**(B)** Representative wide-field images of ovaries from wildtype, *dtorsin*$^{KO13}$ and *dtorsin*$^{KO13}$ + *Rescue*$^{Dup}$ females 4 days after eclosion. Scale bars: 200 µm.

**(C)** Quantification of egg chamber development in ovaries from wild-type, *dtorsin*$^{KO}$ and *dtorsin*$^{KO13}$ + *Rescue*$^{Dup}$ females (N = 3, n = 20 ovarioles).

**(D)** Fertility assay of wildtype, *dtorsin*$^{KO}$ and *dtorsin*$^{KO13}$ + *Rescue*$^{Dup}$ females. Each dot represents the number of progeny from a single female crossed to two wildtype males (N = 3 independent crosses, **** $p \leq 0.0001$, *** $p \leq 0.001$, student's unpaired t-test, two-tailed, mean +/- SD).

**(E)** Quantification of the nurse cell and oocyte number per egg chamber (N = 3, n = 25 egg chambers in stage 4/5, ns $p > 0.05$, unpaired student's t-test, two-tailed).

**(F)** Representative confocal images of stage 4/5 egg chambers from ovaries of the indicated genotypes. Ovaries were fixed and immunostained with mAb414 (anti-FG-Nups, yellow) antibody. DNA was stained with DAPI (magenta). Scale bars: 10 µm.

**(G)** Quantification of the number of FG-Nups-positive clusters per nurse cell nucleus in stage 4/5 egg chambers from (F) using Z-stack confocal images (N = 3, n = 150, **** $p \leq 0.0001$, student's unpaired t-test, two-tailed, mean +/- SD).

**(H)** Quantification of the diameter of FG-Nups-positive perinuclear foci in in stage 4/5 egg chambers from (F) (N = 3, n = 150, mean +/- SD).

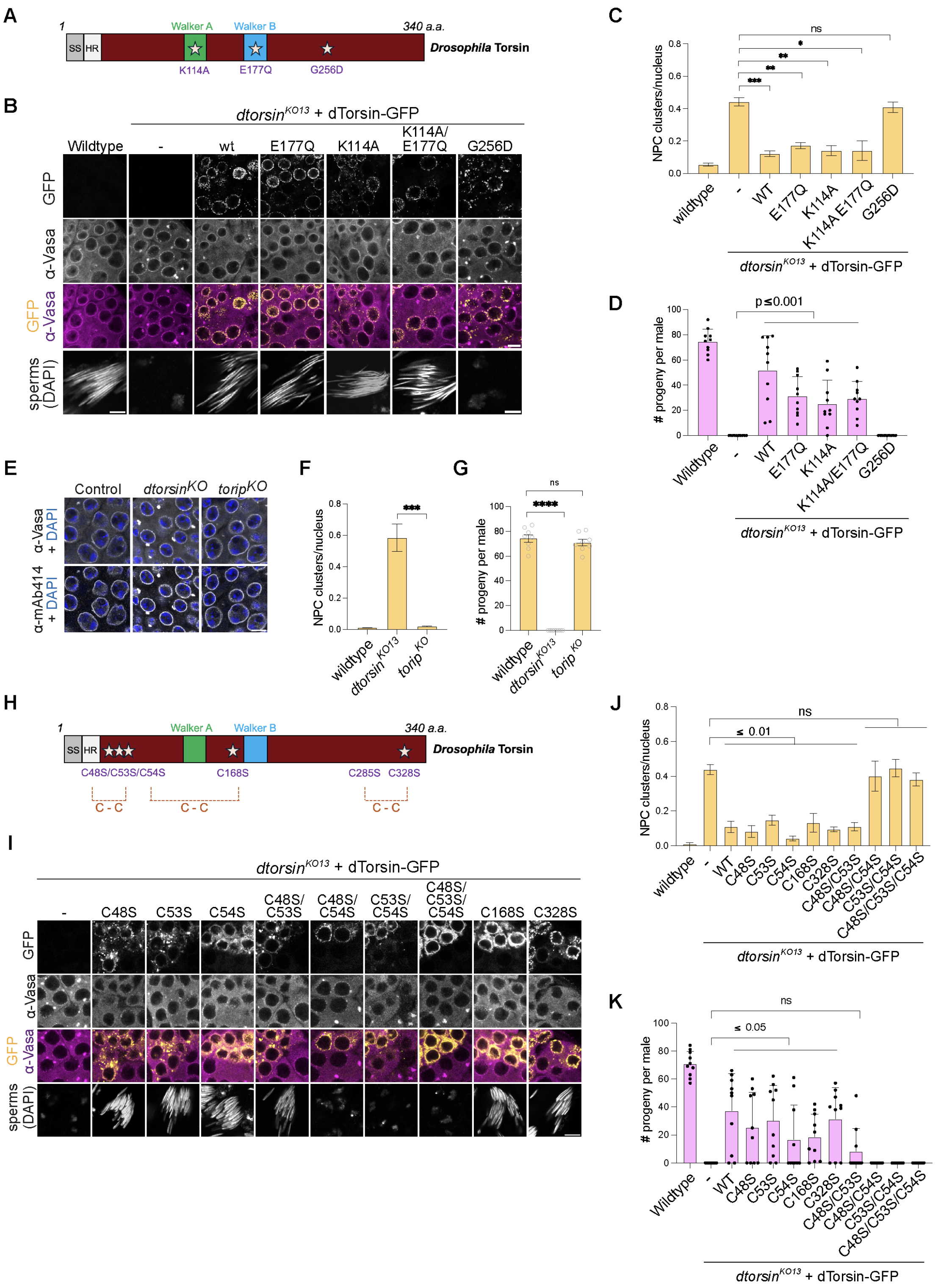


**Figure S4: dTorsin oligomerization and conserved cysteine residues are required for dTorsin's function in spermatogenesis.**

**(A)** Schematic of dTorsin showing the mutated residues (stars) in the Walker A (K114) and Walker B (E177) motifs as well as the G256D mutant affecting the multimerization interface. SS - signal sequence, HR – hydrophobic region, a.a. – amino acids.

**(B)** Representative confocal images of spermatocytes and sperm nuclei bundles from control or *dtorsin*$^{KO}$ testes expressing the indicated GFP-tagged UASt-dTorsin transgenes (yellow) under the control of *bam-Gal4*. Testes were immunostained with anti-Vasa antibodies (magenta) and DNA stained with DAPI (white). Scale bars: 10 µm (spermatocytes), 5 µm (sperm nuclei bundles).

**(C)** Quantification of Vasa-positive perinuclear foci in spermatocytes from (B) (N = 3, n ≥ 250; *** p ≤ 0.001, ** p ≤ 0.01, * p ≤ 0.05, ns p > 0.05; student's unpaired t-test, two-tailed, mean +/- SEM).

**(D)** Fertility assay of males of the indicated genotypes. Each dot represents number of progeny from a single male fly crossed to two virgin females (N = 10 independent crosses; **** p ≤ 0.0001, *** p ≤ 0.001, ** p ≤ 0.01, * p ≤ 0.05, ns p > 0.05; student's unpaired t-test, two-tailed, mean +/- SEM).

**(E)** Representative confocal images of control, *dtorsin*$^{KO}$ and *torip*$^{KO}$ spermatocytes. Testes were fixed and immunostained with mAb414 (anti-FG-Nups) or anti-Vasa antibodies (white). DNA was stained with DAPI (blue). Scale bar: 10 µm.

**(F)** Quantification of number of Vasa/FG-Nups-positive perinuclear foci from (E) (N = 3, n ≥ 240, *** p ≤ 0.001, student's unpaired t-test, two-tailed, mean +/- SEM).

**(G)** Fertility assay of *torip*$^{KO}$ males. Each dot represents progeny number of a single male (N = 8 independent crosses; **** p ≤ 0.0001, ns p > 0.05; student's unpaired t-test, two-tailed, mean +/- SEM).

**(H)** Schematic depiction of dTorsin showing the positions of cysteine residues, labelling as in (A). The dashed lines show putative disulfide bridges.

**(I)** Representative confocal images of *dtorsin*$^{KO}$ testes expressing *UASt-dTorsin-GFP* cysteine mutant transgenes (yellow) under the control of *bam-Gal4*, immunostained with anti-Vasa antibody (magenta). Scale bars: 10 µm (spermatocytes), 5 µm (sperm nuclei bundles).

**(J)** Quantification of Vasa-positive perinuclear foci from (I) (N = 3, n ≥ 220, *** p ≤ 0.001, ** p ≤ 0.01, * p ≤ 0.05, ns p > 0.05, student's unpaired t-test, two-tailed, mean +/- SEM).

**(K)** Fertility assay of males expressing the indicated GFP-tagged dTorsin variants. Each dot represents number of progeny from a single male crossed to two wild-type females (N = 10 independent crosses, *** p ≤ 0.001, ** p ≤ 0.01, * p ≤ 0.05, ns p > 0.05; student's unpaired t-test, two-tailed, mean +/- SEM).

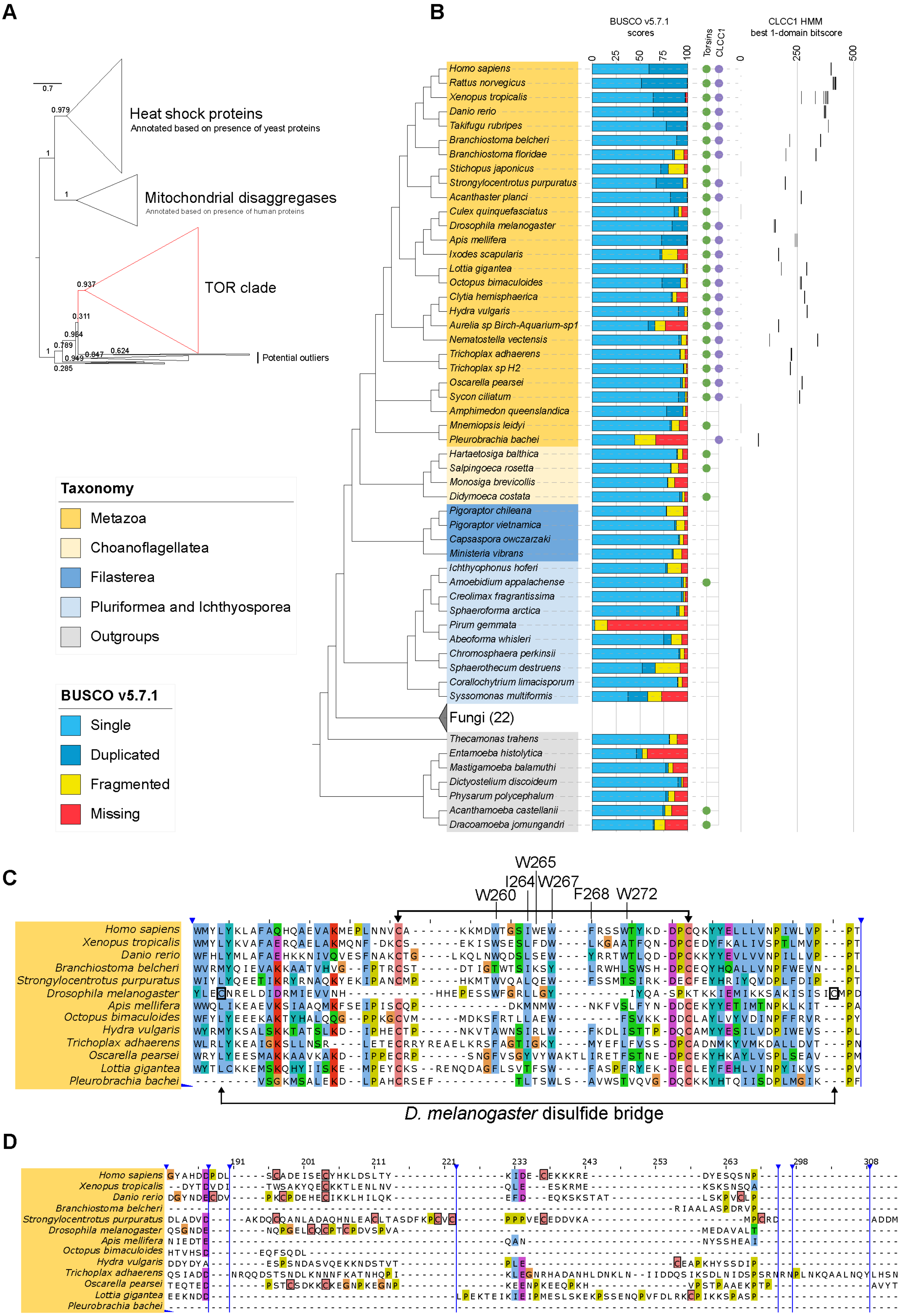
A
0.7
0.979
1
1
Heat shock proteins
Annotated based on presence of yeast proteins
Mitochondrial disaggregases
Annotated based on presence of human proteins
0.937
TOR clade
0.311
0.984
0.789
1
0.949
0.847
0.624
0.285
Potential outliers
Taxonomy
Metazoa
Choanoflagellatea
Filasterea
Pluriformea and Ichthyosporea
Outgroups
BUSCO v5.7.1
Single
Duplicated
Fragmented
Missing
B
BUSCO v5.7.1 scores
0
25
50
75
100
Torsins
CLCC1
CLCC1 HMM best 1-domain bitscore
0
250
500
Homo sapiens
Rattus norvegicus
Xenopus tropicalis
Danio rerio
Takifugu rubripes
Branchiostoma belcheri
Branchiostoma floridae
Stichopus japonicus
Strongylocentrotus purpuratus
Acanthaster planci
Culex quinquefasciatus
Drosophila melanogaster
Apis mellifera
Ixodes scapularis
Lottia gigantea
Octopus bimaculoides
Clytia hemisphaerica
Hydra vulgaris
Aurelia sp Birch-Aquarium-sp1
Nematostella vectensis
Trichoplax adhaerens
Trichoplax sp H2
Oscarella pearsei
Sycon ciliatum
Amphimedon queenslandica
Mnemiopsis leidyi
Pleurobrachia bachei
Hartaetosiga balthica
Salpingoeca rosetta
Monosiga brevicollis
Didymoeca costata
Pigoraptor chileana
Pigoraptor vietnamica
Capsaspora owczarzaki
Ministeria vibrans
Ichthyophonus hoferi
Amoebidium appalachense
Creolimax fragrantissima
Sphaeroforma arctica
Pirum gemmata
Abeoforma whisleri
Chromosphaera perkinsii
Sphaerothecum destruens
Corallochytrium limacisporum
Syssomonas multiformis
Fungi (22)
Thecamonas trahens
Entamoeba histolytica
Mastigamoeba balamuthi
Dictyostelium discoideum
Physarum polycephalum
Acanthamoeba castellanii
Dracoamoeba jomungandri
C
W265
I264
W267
W260
F268
W272
Homo sapiens
Xenopus tropicalis
Danio rerio
Branchiostoma belcheri
Strongylocentrotus purpuratus
Drosophila melanogaster
Apis mellifera
Octopus bimaculoides
Hydra vulgaris
Trichoplax adhaerens
Oscarella pearsei
Lottia gigantea
Pleurobrachia bachei
D. melanogaster disulfide bridge
D
191
201
211
221
233
243
253
263
298
308
Homo sapiens
Xenopus tropicalis
Danio rerio
Branchiostoma belcheri
Strongylocentrotus purpuratus
Drosophila melanogaster
Apis mellifera
Octopus bimaculoides
Hydra vulgaris
Trichoplax adhaerens
Oscarella pearsei
Lottia gigantea
Pleurobrachia bachei

**Figure S5: Figure legend: Phylogenetic profiles of the Torsin family and CLCC1 family across Opisthokonta.**

**(A)** Gene tree of the Torsin family and other closely related protein families as labelled in the panel. The monophyletic clade containing the known human Torsins is highlighted in red and was used to build the phylogenetic profile. The gene tree was constructed using FastTree. The numbers against different branches are bootstrap supports (closer to 1, better the score). The tree scale is in substitutions per site.

**(B)** Distribution of Torsins and CLCC1 plotted against a taxonomic species tree. Species names are coloured based on taxonomy. Proteomes for each species were assessed using BUSCO v5.7.1 with the eukaryota lineage of 255 marker proteins. The scale of the BUSCO graph is in percent. Filled circles in the profile indicate that at least one ortholog is present. The scale of the CLCC1 HMM best 1-domain scores graph is unitless.

**(C)** Multiple sequence alignment of the CLCC1 family showing the amphipathic helix region. This alignment is a subset of the CLCC1 alignment used to build the phylogenetic profile in (B). Species were selected to represent the known diversity of metazoa, and to include species of interest. Arrows at the top of the alignment indicate the cysteines forming the loop disulfide bridge in mammals, arrows at the bottom indicate the position of the respective cysteines in *D. melanogaster*. Hydrophobic residues contained in amphipathic helix 1 (AH1) in human CLCC1 are highlighted above. Vertical blue lines indicate positions where columns composed exclusively of gaps were hidden.

**(D)** Multiple sequence alignment of the CLCC1 family showing the N-terminal region. This alignment is a subset of the CLCC1 alignment used to build the phylogenetic profile in (B). Species were selected to represent the known diversity of Metazoa, and to include species of interest. Cysteines in this region are manually annotated and are indicated by a pink box with a solid, black border. Vertical blue lines indicate positions where columns composed exclusively of gaps were hidden.

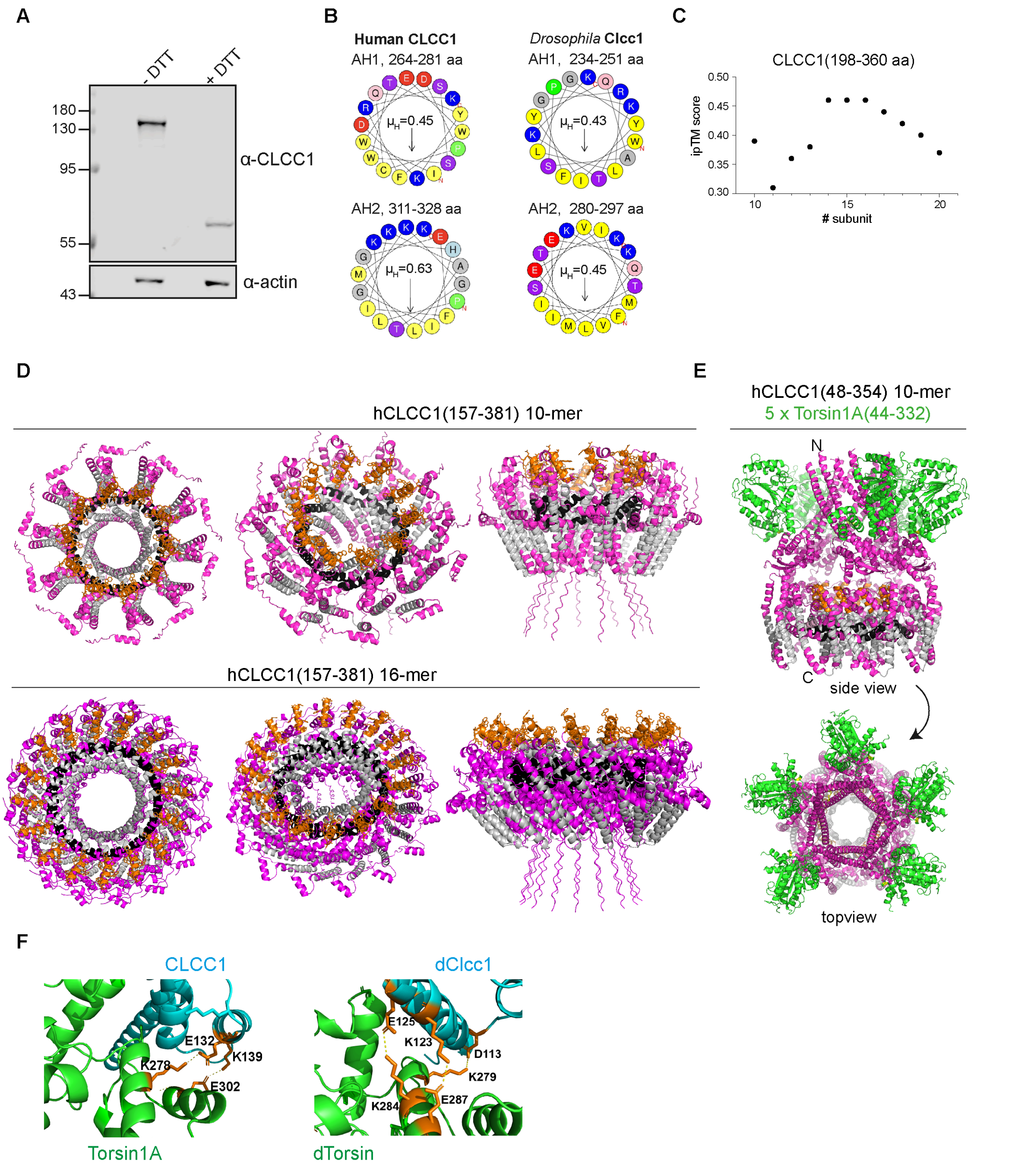


**Figure S6: CLCC1 as disulfide-linked dimer, its amphipathic helices AH1 and AH2 and the ring-shaped CLCC1 10- and 16-mers**

**(A)** Immunoblotting of HeLa cells detecting endogenous CLCC1 in presence (reducing condition) or absence (non-reducing condition) of 50 mM DTT in the SDS lysis buffer.

**(B)** Helical wheel projection of amphipathic helices AH1 and AH2 of human and *Drosophila* CLCC1, showing the hydrophobic moment ($\mu_H$) calculated using HeliQuest (Gautier et al., 2008).

**(C)** Plot of ipTM scores of higher-order CLCC1 assemblies (aa 198 – 360) based on AlphaFold-Multimer. Oligomers with fewer than 10 subunits seem insufficient to form a ring-ling structure.

**(D)** AlphaFold Multimer prediction of ring-shaped multimer (16-mer) of human CLCC1(157-381) (pink), showing amphipathic helix AH1 (orange) protruding outside of the structure as well as AH2 (black) embedded into the oligomer intercalated by transmembrane (TM) regions (grey).

**(E)** AlphaFold Multimer prediction of ring-shaped multimer (10-mer) of human CLCC1(48-354) together with 5 Torsin1A molecules (44-332).

**(F)** Zoom representation of the interaction interfaces, with key interacting residues that form salt bridges labeled and in stick representation, based on the CLCC1-Torsin1A complex (left) predicted by (Zhang et al., 2025) and dTorsin-dCLCC1 model (right), based on a structure predicted by AFM. For the dTorsin interface mutant, K279E, K284E, E287K (original residues in orange) were mutated to disrupt dTorsin-dCLCC1 interaction (Fig. 3).

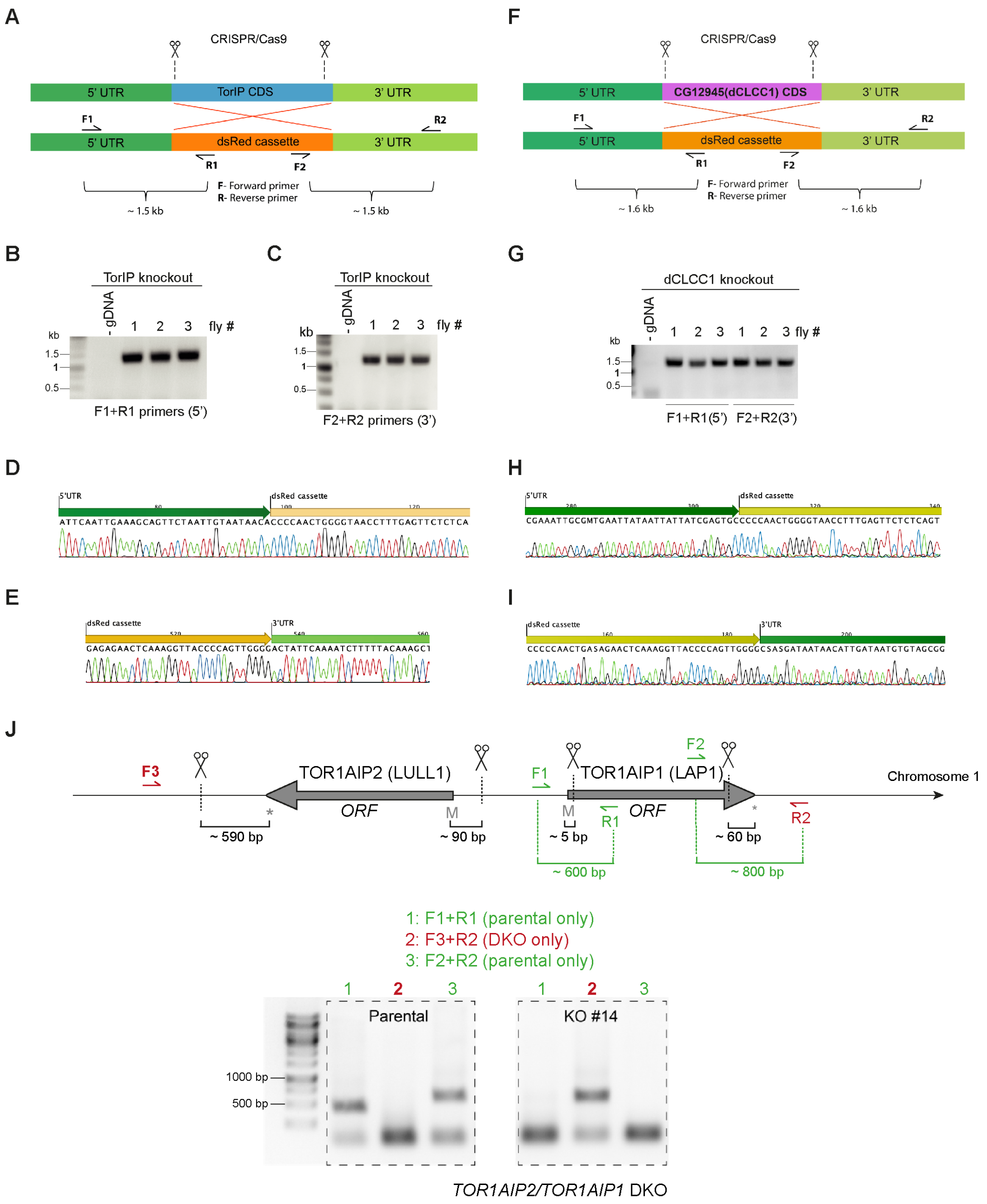


**Figure S7: Characterization of *torip* and *Drosophila clcc1* knockout strains.**

**(A)** Schematic of *Torip* knockout generation. The integration cassette contains a 3xP3-driven DsRed cassette for selection. The sequences used for cloning homology arms (5' and 3') as well as primer positions for strain characterization by PCR are shown.

**(B, C, D, E)** PCR on genomic DNA of three homozygous *torip*$^{KO}$ male flies using the indicated primers. Bands were cut out of the gel, purified and sent for Sanger sequencing (D: F1+R1, E: F2+R2).

**(F)** Schematic of *dClcc1* knockout generation. The integration cassette contains a 3xP3-driven DsRed cassette for selection. The sequences used for cloning homology arms (5' and 3') as well as primer positions for strain characterization by PCR are shown.

**(G, H, I)** PCR on genomic DNA of three homozygous *dclcc1*$^{KO}$ male flies using the indicated primers. Bands were cut out of the gel, purified and sent for Sanger sequencing.

**(J)** Characterization of the *HeLa TOR1AIP1/TOR1AIP2* double knockout (DKO) cell line by PCR. The forward primer F3 was designed ~200 bp downstream of the TOR1AIP2 3' Cas9 cutting site, and the reverse primer R2 was positioned ~200 bp downstream of the 3’ Cas9 cutting site of the TOR1AIP1. This primer pair produced a clear amplification band in the DKO cell line but not in the parental HeLa line, consistent with the expected genomic deletion (the distance between F3 and R2 in the parental genome is ~73.5 kb). As internal controls, PCRs targeting the TOR1AIP1 Cas9 cutting sites were performed using primer pairs F1/R1 (5′ cut site; expected amplicon ~600 bp) and F2/R2 (3′ cut site; expected amplicon ~800 bp).

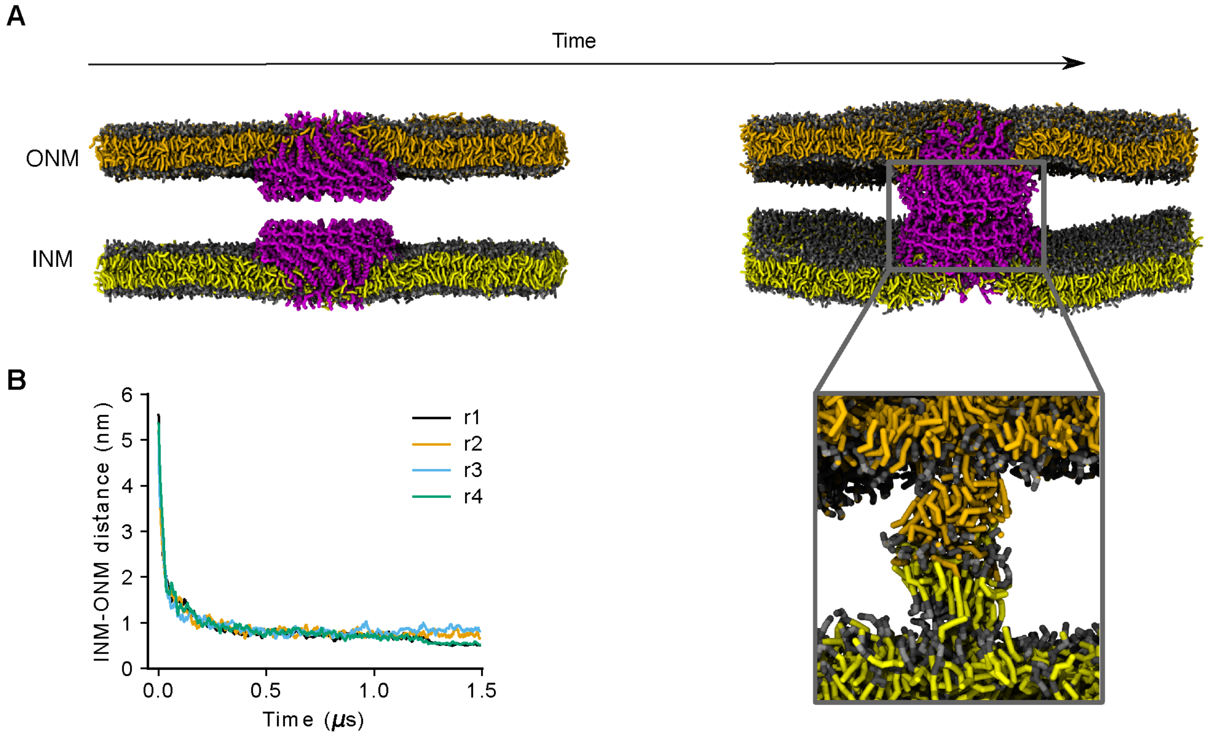


**Figure S8: Molecular dynamics simulations of membrane fusion mediated by AH1 helices of two opposing CLCC1 16-mers.**

**(A)** Coarse-grained MD simulation snapshots of a two planar bilayer opposed to each other with CLCC1 16-mer inserted in both, akin to what has been suggested for yeast BRL1-like proteins (Fischer et al., 2025). Inset shows lipid mixing inside the channel formed by both CLCC1 rings.

**(B)** Time-trace of minimum INM-ONM distance during the simulation showing stable contact and mixing between lipids of opposing membranes in (D).

**Supplementary Table 1: Protein enriched on Torsin1A by TurboID**

Proteins identified by mass spectrometry that were significantly enriched in the Torsin1A sample compared to the GFP control by *in vivo* proximity labelling in HeLa cells. Proteins with adjusted p-values < 0.01 and $log_2$-fold enrichment > 2 are listed, sorted by fold change. Putative ER-resident factors are highlighted in bold. Torsin activators LAP1 and LULL1 in orange, CLCC1 in violet.

| **ID** | **Gene name** | **log2FC (Torsin1A vs GFP)** | **p-value** |
|---|---|---|---|
| **O14656** | **TOR1A** | **11.79348** | **0.000148** |
| Q96B26 | EXOSC8 | 6.194858 | 0.001941 |
| Q9Y3D8 | AK6 | 5.949323 | 0.002594 |
| **Q96HD1** | **CRELD1** | **5.707139** | **0.004416** |
| **Q8TBM8** | **DNAJB14** | **5.526229** | **0.001758** |
| **Q9UHQ4** | **BCAP29** | **5.426767** | **0.001286** |
| Q9NX62 | IMPAD1 | 5.055913 | 0.00628 |
| **Q9BXS4** | **TMEM59** | **4.947359** | **0.008057** |
| **Q9NPA0** | **EMC7** | **4.834972** | **0.004663** |
| P17252 | PRKCA | 4.677055 | 0.004296 |
| **P04035** | **HMGCR** | **4.547536** | **0.001999** |
| Q658P3 | STEAP3 | 4.523902 | 0.001769 |
| Q9P2E5 | CHPF2 | 4.509613 | 0.007292 |
| **Q96S66** | **CLCC1** | **4.496696** | **0.001817** |
| **Q7Z388** | **DPY19L4** | **4.495312** | **0.002161** |
| Q969F9 | HPS3 | 4.477615 | 0.007808 |
| **O75460** | **ERN1** | **4.429972** | **0.002611** |
| **P62341** | **SELENOT** | **4.386349** | **0.008804** |
| Q8TBC4 | UBA3 | 4.212486 | 0.000569 |
| O00471 | EXOC5 | 4.171322 | 0.001443 |
| **Q8NC56** | **LEMD2** | **4.09539** | **0.00258** |
| O43264 | ZW10 | 4.088616 | 0.00003 |
| P51659 | HSD17B4 | 4.070062 | 0.009996 |
| P43357 | MAGEA3 | 3.984806 | 0.003003 |
| O94919 | ENDOD1 | 3.974392 | 0.00295 |
| **Q6PKC3** | **TXNDC11** | **3.846837** | **0.001082** |
| P08069 | IGF1R | 3.794462 | 0.008822 |
| Q96EH3 | MALSU1 | 3.737491 | 0.006567 |
| P25445 | FAS | 3.736696 | 0.003389 |
| **Q7L5N7** | **LPCAT2** | **3.728131** | **0.003389** |
| **Q5JTV8** | **LAP1** | **3.713171** | **0.007776** |
| **O60353** | **FZD6** | **3.693651** | **0.005712** |
| O95466 | FMNL1 | 3.646003 | 0.002779 |
| Q9HCG8 | CWC22 | 3.629838 | 0.001852 |
| Q9H4L7 | SMARCAD1 | 3.596229 | 0.003496 |
| **Q9H497** | **TOR3A** | **3.552268** | **0.009739** |
| O15431 | SLC31A1 | 3.458982 | 0.009236 |
| **Q9GZP9** | **DERL2** | **3.44288** | **0.008953** |

| | | | |
|---|---|---|---|
| O96017 | CHEK2 | 3.438927 | 0.004417 |
| Q8IYB7 | DIS3L2 | 3.435019 | 0.004416 |
| O14763 | TNFRSF10B | 3.362124 | 0.001778 |
| P52788 | SMS | 3.332464 | 0.00941 |
| **Q8N766** | **EMC1** | **3.331213** | **0.009739** |
| P98153 | DGCR2 | 3.298424 | 0.00535 |
| O95168 | NDUFB4 | 3.265703 | 0.00123 |
| Q9Y287 | ITM2B | 3.230992 | 0.001758 |
| Q9Y221 | NIP7 | 3.228203 | 0.007849 |
| Q9BXI9 | C1QTNF6 | 3.188007 | 0.008057 |
| P18074 | ERCC2 | 3.185534 | 0.002301 |
| Q9BRQ8 | AIFM2 | 3.170085 | 0.003496 |
| Q9H0P0 | NT5C3A | 3.11535 | 0.002691 |
| Q99808 | SLC29A1 | 3.084801 | 0.008758 |
| Q14651 | PLS1 | 3.032474 | 0.007012 |
| **Q96HV5** | **TMEM41A** | **2.944689** | **0.00295** |
| **Q5U4P2** | **ASPHD1** | **2.891801** | **0.001852** |
| **Q8NFQ8** | **LULL1** | **2.865674** | **0.00759** |
| **Q99942** | **RNF5** | **2.830726** | **0.003389** |
| P30740 | SERPINB1 | 2.813969 | 0.001769 |
| O14828 | SCAMP3 | 2.694643 | 0.006009 |
| Q92600 | CNOT9 | 2.669709 | 0.004416 |
| P05067 | APP | 2.623083 | 0.001815 |
| **O15533** | **TAPBP** | **2.581031** | **0.004591** |
| Q9BV23 | ABHD6 | 2.395965 | 0.008498 |
| Q86SF2 | GALNT7 | 2.389394 | 0.00941 |
| Q8NBI5 | SLC43A3 | 2.308217 | 0.009988 |
| Q92499 | DDX1 | 2.261496 | 0.000974 |
| **Q6PJF5** | **RHBDF2** | **2.216907** | **0.002301** |
| Q96BW9 | TAMM41 | 2.209716 | 0.003247 |
| Q9HCM3 | KIAA1549 | 2.106466 | 0.008953 |
| **Q8NI22** | **MCFD2** | **2.09618** | **0.009856** |
| Q9H6R4 | NOL6 | 2.093602 | 0.006174 |
| P40925 | MDH1 | 2.05515 | 0.005772 |

**Supplementary Table 2: Metazoa proteomes taken from UniProt**

| **Species name** | **Abbreviation** |
|---|---|
| *Xenopus tropicalis* | Chor0 |
| *Ailuropoda melanoleuca* | Chor1 |
| *Chrysochloris asiatica* | Chor2 |
| *Danio rerio* | Chor3 |
| *Enhydra lutris kenyoni* | Chor4 |
| *Equus caballus* | Chor5 |
| *Felis catus* | Chor6 |
| *Homo sapiens* | Chor7 |
| *Macaca mulatta* | Chor8 |
| *Mus musculus* | Chor9 |
| *Neomonachus schauinslandi* | Chor10 |
| *Oryzias latipes* | Chor11 |
| *Phascolarctos cinereus* | Chor12 |
| *Phyllostomus discolor* | Chor13 |
| *Pogona vitticeps* | Chor14 |
| *Rattus norvegicus* | Chor15 |
| *Xenopus laevis* | Chor16 |
| *Apteryx mantelli mantelli* | Chor17 |
| *Otolemur garnettii* | Chor18 |
| *Loxodonta africana* | Chor19 |
| *Oikopleura dioica* | Chor20 |
| *Branchiostoma floridae* | Chor21 |
| *Anolis carolinensis* | Chor22 |
| *Ornithorhynchus anatinus* | Chor23 |
| *Takifugu rubripes* | Chor24 |
| *Eublepharis macularius* | Chor25 |
| *Strongylocentrotus purpuratus* | Chor26 |
| *Ciona intestinalis* | Chor27 |
| *Latimeria chalumnae* | Chor28 |
| *Tupaia chinensis* | Chor29 |
| *Scleropages formosus* | Chor30 |
| *Carlito syrichta* | Chor31 |
| *Aquarana catesbeiana* | Chor32 |
| *Stichopus japonicus* | Chor33 |
| *Chiloscyllium punctatum* | Chor34 |
| *Scyliorhinus torazame* | Chor35 |

| | |
|---|---|
| *Callorhinchus milii* | Chor36 |
| *Vombatus ursinus* | Chor37 |
| *Suricata suricatta* | Chor38 |
| *Notechis scutatus* | Chor39 |
| *Panthera pardus* | Chor40 |
| *Branchiostoma belcheri* | Chor41 |
| *Sphenodon punctatus* | Chor42 |
| *Prolemur simus* | Chor43 |
| *Salvator merianae* | Chor44 |
| *Dromaius novaehollandiae* | Chor45 |
| *Gallus gallus* | Chor46 |
| *Taeniopygia guttata* | Chor47 |
| *Alligator sinensis* | Chor48 |
| *Crocodylus porosus* | Chor49 |
| *Pelusios castaneus* | Chor50 |
| *Chrysemys picta bellii* | Chor51 |
| *Gopherus evgoodei* | Chor52 |
| *Varanus komodoensis* | Chor53 |
| *Rousettus aegyptiacus* | Chor54 |
| *Leptobrachium leishanense* | Chor55 |
| *Erpetoichthys calabaricus* | Chor56 |
| *Acanthaster planci* | Chor57 |
| *Python bivittatus* | Chor58 |
| *Galemys pyrenaicus* | Chor59 |
| *Eleutherodactylus coqui* | Chor60 |
| *Hymenochirus boettgeri* | Chor61 |
| *Albula goreensis* | Chor62 |
| *Branchiostoma lanceolatum* | Chor63 |
| *Anguilla anguilla* | Chor64 |
| *Megalops atlanticus* | Chor65 |
| *Synaphobranchus kaupii* | Chor66 |
| *Podarcis lilfordi* | Chor67 |
| *Sciurus vulgaris* | Chor68 |
| *Aldrovandia affinis* | Chor69 |
| *Acipenser oxyrinchus oxyrinchus* | Chor70 |
| *Pelobates cultripes* | Chor71 |
| *Petromyzon marinus* | Chor72 |
| *Crotalus adamanteus* | Chor73 |

| | |
|---|---|
| *Pleurodeles waltl* | Chor75 |
| *Gnathostoma spinigerum* | Chor76 |
| *Ditylenchus destructor* | Chor77 |
| *Steinernema hermaphroditum* | Chor78 |
| *Geodia barretti* | Chor79 |
| *Paramuricea clavata* | Chor80 |
| *Parascaris univalens* | Chor81 |
| *Acrobeloides nanus* | Chor82 |
| *Drosophila melanogaster* | Chor83 |
| *Brachionus calyciflorus* | Chor84 |
| *Bursaphelenchus xylophilus* | Chor85 |
| *Clytia hemisphaerica* | Chor86 |
| *Meloidogyne enterolobii* | Chor87 |
| *Actinia tenebrosa* | Chor88 |
| *Panagrellus redivivus* | Chor89 |
| *Brachionus plicatilis* | Chor90 |
| *Dracunculus medinensis* | Chor91 |
| *Enterobius vermicularis* | Chor92 |
| *Trichoplax sp. H2* | Chor93 |
| *Hypsibius exemplaris* | Chor94 |
| *Ramazzottius varieornatus* | Chor95 |
| *Trichinella nativa* | Chor96 |
| *Haemonchus contortus* | Chor97 |
| *Trichoplax adhaerens* | Chor98 |
| *Amphimedon queenslandica* | Chor99 |
| *Brugia malayi* | Chor100 |
| *Pristionchus pacificus* | Chor101 |
| *Caenorhabditis elegans* | Chor102 |
| *Nematostella vectensis* | Chor103 |
| *Anopheles darlingi* | Chor104 |
| *Magallana gigas* | Chor105 |
| *Schistosoma mansoni* | Chor106 |
| *Strigamia maritima* | Chor107 |
| *Helobdella robusta* | Chor108 |
| *Echinococcus multilocularis* | Chor109 |
| *Lottia gigantea* | Chor110 |
| *Macrostomum lignano* | Chor111 |
| *Hymenolepis diminuta* | Chor112 |

| | |
|---|---|
| *Opisthorchis felineus* | Chor113 |
| *Octopus vulgaris* | Chor114 |
| *Varroa destructor* | Chor115 |
| *Crassostrea virginica* | Chor116 |
| *Owenia fusiformis* | Chor117 |
| *Daphnia sinensis* | Chor118 |
| *Biomphalaria glabrata* | Chor119 |
| *Nephila pilipes* | Chor120 |
| *Cichlidogyrus casuarinus* | Chor122 |

**Supplementary Table 3: 74 Opisthokont proteomes**

| Name | Abbreviation | Source |
|---|---|---|
| *Abeoforma whisleri* | Awhi | https://doi.org/10.7554/eLife.26036 |
| *Acanthamoeba castellanii* | Acas | EukProt |
| *Acanthaster planci* | Apla | UniProt |
| *Allomyces macrogynus* | Amac | EukProt |
| *Amoebidium appalachense* | Aapp | https://doi.org/10.1126/sciadv.ado6406 |
| *Amphimedon queenslandica* | Aque | EukProt |
| *Apis mellifera* | Amel | EukProt |
| *Aspergillus nidulans* | Anid | EukProt |
| *Aurelia sp Birch-Aquarium-sp1* | Asp | EukProt |
| *Batrachochytrium dendrobatidis* | Bden | EukProt |
| *Branchiostoma belcheri* | Bbel | UniProt |
| *Branchiostoma floridae* | Bflo | EukProt |
| *Candida albicans* | Calb | EukProt |
| *Capsaspora owczarzaki* | Cowc | EukProt |
| *Chromosphaera perkinsii* | Cper | https://doi.org/10.7554/eLife.26036 |
| *Clytia hemisphaerica* | Chem | EukProt |
| *Coprinopsis cinerea* | Ccin | EukProt |
| *Corallochytrium limacisporum* | Clim | EukProt |
| *Creolimax fragrantissima* | Cfra | https://doi.org/10.7554/eLife.08904 |
| *Cryptococcus gattii* | Cgat | EukProt |
| *Cryptococcus neoformans* | Cneo | EukProt |
| *Culex quinquefasciatus* | Cqui | EukProt |
| *Danio rerio* | Drer | UniProt |
| *Dictyostelium discoideum* | Ddis | EukProt |
| *Didymoeca costata* | Dcos | EukProt |
| *Dracoamoeba jomungandri* | Djom | EukProt |
| *Entamoeba histolytica* | Ehis | EukProt |
| *Rhizophagus irregularis* | Rirr | EukProt |
| *Rhizopus delemar* | Rdel | EukProt |
| *Lobosporangium transversale* | Ltra | EukProt |
| *Basidiobolus meristosporus* | Bmer | EukProt |
| *Coemansia reversa* | Crev | EukProt |
| *Fonticula alba* | Falb | EukProt |
| *Gonapodya prolifera* | Gpro | EukProt |
| *Hartaetosiga balthica* | Hbal | EukProt |
| *Hydra vulgaris* | Hvul | UniProt |
| *Ichthyophonus hoferi* | Ihof | https://www.sciencedirect.com/science/article/pii/S0960982215008878 |
| *Ixodes scapularis* | Isca | EukProt |
| *Laccaria bicolor* | Lbic | EukProt |
| *Lottia gigantea* | Lgig | EukProt |
| *Mastigamoeba balamuthi* | Mbal | EukProt |
| *Ministeria vibrans* | Mvib | https://doi.org/10.1038/s41586-022-05110-4, https://doi.org/10.6084/m9.figshare.13140191.v1 |
| *Mnemiopsis leidyi* | Mlei | EukProt |
| *Monosiga brevicollis* | Mbre | UniProt |

| | | |
|---|---|---|
| *Nematostella vectensis* | Nvec | EukProt |
| *Neurospora crassa* | Ncra | EukProt |
| *Octopus bimaculoides* | Obim | EukProt |
| *Oscarella pearsei* | Opea | EukProt |
| *Parvularia atlantis* | Patl | https://doi.org/10.1038/s41586-022-05110-4, https://doi.org/10.6084/m9.figshare.13140191.v1 |
| *Physarum polycephalum* | Ppol | EukProt |
| *Pigoraptor chileana* | Pchi | https://doi.org/10.1038/s41586-022-05110-4, https://doi.org/10.6084/m9.figshare.13140191.v1 |
| *Pigoraptor vietnamica* | Pvie | https://doi.org/10.1038/s41586-022-05110-4, https://doi.org/10.6084/m9.figshare.13140191.v1 |
| *Pirum gemmata* | Pgem | https://doi.org/10.7554/eLife.26036 |
| *Pleurobrachia bachei* | Pbac | EukProt |
| *Rattus norvegicus* | Rnor | EukProt |
| *Rozella allomycis* | Rall | EukProt |
| *Saccharomyces cerevisiae* | Scer | UniProt |
| *Salpingoeca rosetta* | Sros | EukProt |
| *Schizosaccharomyces pombe* | Spom | UniProt |
| *Sphaeroforma arctica* | Sarc4 | https://doi.org/10.7554/eLife.49801 |
| *Sphaerothecum destruens* | Sdes | https://doi.org/10.1038/s41586-022-05110-4 |
| *Spizellomyces punctatus* | Spun | EukProt |
| *Stichopus japonicus* | Sjap | UniProt |
| *Strongylocentrotus purpuratus* | Spur | EukProt |
| *Sycon ciliatum* | Scil | EukProt |
| *Syssomonas multiformis* | Smul | EukProt |
| *Takifugu rubripes* | Trub | EukProt |
| *Thecamonas trahens* | Ttra | EukProt |
| *Trichoplax adhaerens* | Tadh | EukProt |
| *Trichoplax sp H2* | Tsp | EukProt |
| *Drosophila melanogaster* | Dmel | UniProt |
| *Homo sapiens* | Hsap | UniProt |
| *Ustilago maydis 521* | Umay | EukProt |
| *Xenopus tropicalis* | Xtro | EukProt |

### Supplementary Movies

**Movie S1: 3D reconstruction of EGFP-Nup107 fluorescence of spermatocytes from control testes.**

Representative 3D reconstruction of spermatocyte nuclei from fixed testes of control males, expressing EGFP-Nup107, generated using Imaris. Scale bar: 5 μm.

**Movie S2: 3D reconstruction of EGFP-Nup107 fluorescence of spermatocytes from *dtorsin*$^{KO13}$ testes.**

Representative 3D reconstruction of spermatocyte nuclei from fixed testes of *dtorsin*$^{KO13}$ males, expressing EGFP-Nup107, generated using Imaris. Scale bar: 5 μm.
